# The *Toxoplasma gondii* mitochondrial transporter ABCB7 is essential for cytosolic iron-sulfur cluster biogenesis and protein translation

**DOI:** 10.1101/2024.03.15.585200

**Authors:** Andrew E. Maclean, Megan A. Sloan, Eléa A. Renaud, Vincent Demolombe, Sébastien Besteiro, Lilach Sheiner

## Abstract

Iron-sulfur (Fe-S) clusters are ubiquitous inorganic cofactors required for numerous essential cellular pathways. Since they cannot be scavenged from the environment, Fe-S clusters are synthesised de novo in cellular compartments such as the apicoplast, mitochondrion and cytosol. The cytosolic Fe-S cluster biosynthesis pathway relies on transport of an intermediate from the mitochondrial pathway. An ATP binding cassette (ABC) transporter called ABCB7 is responsible for this role in numerous commonly studied organisms, but its role in the medically important apicomplexan parasites has not yet been studied. Here we identify and characterise the *Toxoplasma gondii* ABCB7 homolog. Genetic depletion shows that it is essential for parasite growth, and that disruption triggers partial stage conversion. Characterisation of the knock-down line highlights a defect in cytosolic Fe-S cluster biogenesis leading to defects in protein translation and other pathways including DNA and RNA replication and metabolism. Our work provides support for a broad conservation of the connection between mitochondrial and cytosolic in Fe-S cluster biosynthesis and reveal its importance for parasite survival.

## Introduction

Iron-sulfur (Fe-S) clusters are ubiquitous inorganic metallocofactors, made up of geometric arrangements of iron and sulfur atoms, that are essential across cellular life. Due to their electron transfer capabilities, they are important components of enzymes required for critical cellular functions (1). Many mitochondrial Fe-S proteins are components of the mitochondrial electron transport chain (mETC) and are involved in respiration. Cytosolic and nuclear Fe-S proteins include enzymes that play a role in DNA replication and repair, amino acid biosynthesis, tRNA modification and protein translation (2). Therefore, it is essential for cellular life that Fe-S clusters are synthesised and trafficked to their correct cellular location.

Fe-S clusters cannot be scavenged from the environment, nor from a host cell in the case of intracellular parasites. Therefore, all cells contain dedicated biosynthetic pathways. These directly involve over thirty protein components localised to numerous cell compartments. These proteins are required to bring together and assemble sulfide ions (S^2-^) and ferrous (Fe^2+^) or ferric (Fe^3+^) iron into nascent Fe-S clusters, and then deliver them to numerous client proteins (3). Organisms that contain an endosymbiotic plastid typically contain three such biosynthetic pathways: the mitochondrial Iron-Sulfur Cluster assembly pathway (the ISC pathway), the plastid Sulfur mobilisation pathway (the SUF pathway), and the Cytosolic Iron-sulfur Assembly pathway (the CIA pathway). Whilst the SUF pathway provides Fe-S clusters exclusively for a handful of plastid enzymes, the mitochondrial ISC and cytosolic CIA pathway are connected (Fig. 1A). A mitochondrial cysteine desulfurase (NFS1) liberates sulfur from cysteine, which is then combined with iron and assembled into Fe-S clusters on the mitochondrial scaffold protein ISU1. A number of extra “carrier” proteins then deliver and insert the nascent cluster into mitochondrial

**Fig 1.**
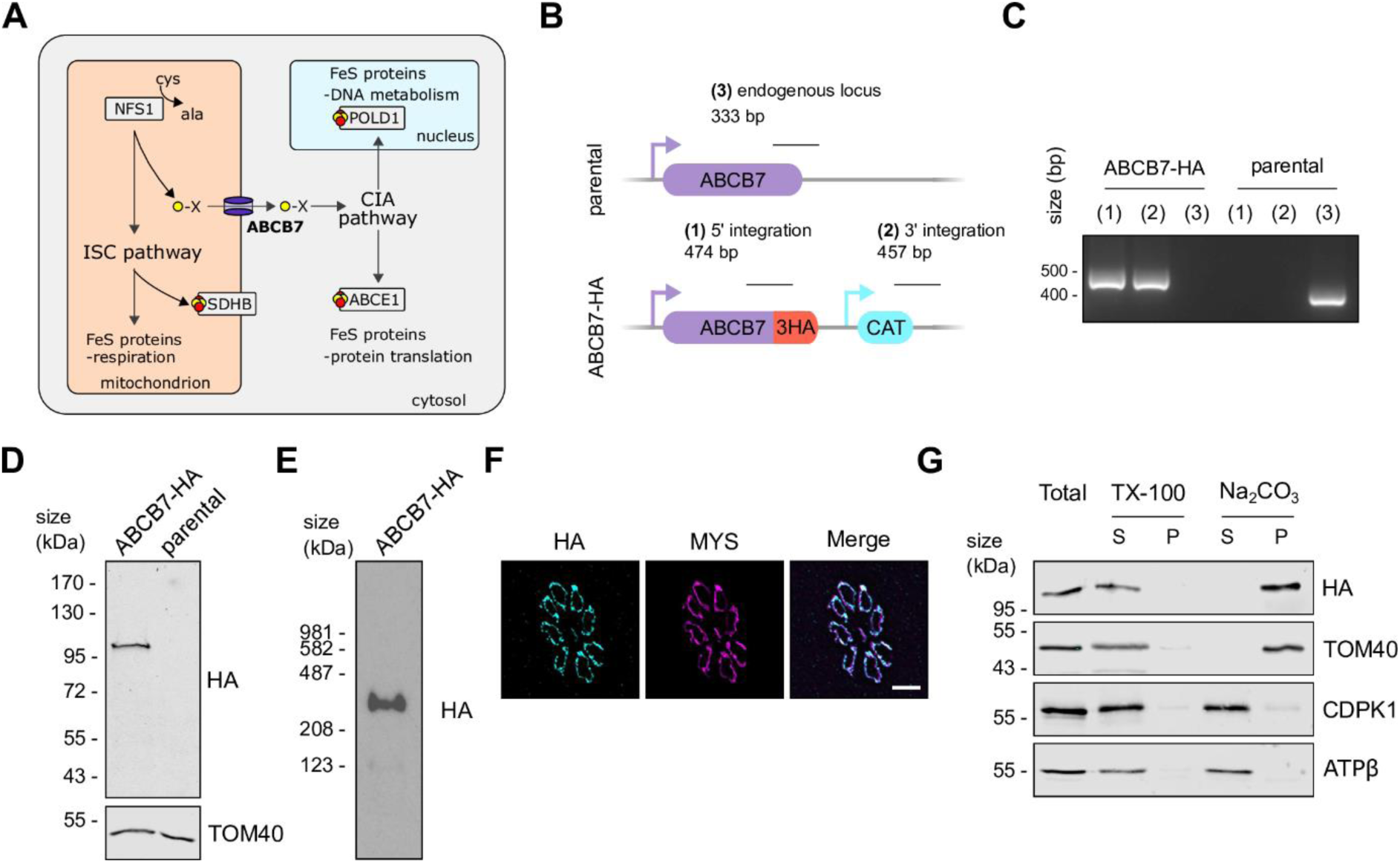
The *Toxoplasma* ABCB7 homolog is an integral membrane protein that localises to the mitochondrion. (A) Schematic of the mitochondrial ISC and cytosolic CIA Fe-S assembly pathways. ABCB7 (bold) connects these two pathways. Fe-S containing proteins evaluated in this study are shown (POLD1, ABCE1 and SDHB) (B) Schematic of the strategy used to C-terminally HA-epitope tag the ABCB7 protein. The expected size of integration PCRs are shown. (C) PCR to test integration of HA-epitope tag and CAT selection cassette into the endogenous locus, as outlined in *(B)*. (D) Immunoblot analysis of whole cell lysate extracted from ABCB7-HA and parental parasites. Samples were separated by SDS-PAGE, blotted, and detected using anti-HA, to visualise ABCB7-HA, and anti-TOM40 as a loading control. (E) BN-PAGE analysis of ABCB7-HA parasites extracted in 1% βDDM, immunolabelled with anti-HA. (F) Immunofluorescence assay analysis of ABCB7-HA parasites, labelled with anti-HA to detect ABCB7-HA (cyan), showing co-localisation with the mitochondrial marker protein MYS (magenta). Scale bar is 5 µM. (G) Immunoblot analysis of ABCB7-HA parasites treated with 1% Triton X-100 (TX-100) or sodium carbonate (Na_2_CO_3_) at pH 11.5 and separated by centrifugation to give a supernatant (S) and pellet (P) fraction. A total fraction, before treatment, (T) is also analysed. ABCB7-HA is found in the pellet fraction of Na_2_CO_3_, like the integral outer membrane protein TOM40. The peripheral mitochondrial matrix subunit of ATP synthase, ATPβ, and the cytosolic protein CDPK1 is found in the soluble supernatant fraction.

Fe-S proteins. The CIA pathway is also dependent on mitochondrial NFS1 (2, 4). A sulfur-containing intermediate produced by NFS1 is exported from the mitochondrion to the cytosol, before being combined with iron and assembled on a cytosolic scaffold protein complex, which contains NBP35 (4). This is followed by a number of extra carrier proteins that help insert the nascent cluster into Fe-S proteins in the cytosol and nucleus (Fig. 1A) called Nar1, CIA1, CIA2 and MMS19. Thus, the cytosolic CIA pathway requires mitochondrial export of sulfur-containing intermediate to function.

Mitochondrial export of this sulfur-containing intermediate is performed by an ATP binding cassette (ABC) transporter called ABCB7 (called ATM1 in yeast and ATM3 in plants), in a glutathione dependent manner (4, 5). ABCB7 is a type IV ABC transporter (6) localised to the inner mitochondrial membrane, with a nucleotide binding domain (NBD) facing the mitochondrial matrix, suggesting it functions as an exporter. ABCB7 transporters operate as half-transporters, with six transmembrane domains from each monomer contributing to forming a channel (7). Initial work in yeast identified ATM1 as the exporter required for cytosolic Fe-S cluster maturation (8). Subsequent studies in yeast, plants and vertebrate models have shown a conserved function, where depletion of this ABC transporter leads to defects in cytosolic and nuclear Fe-S proteins, while the mitochondrial Fe-S proteins remain largely unaffected (9, 10). Recent structural studies in yeasts, plants and mammals have helped elucidate the transport mechanism (11–15), although the exact physiological substrate of the transporter is still the subject of debate. Studies in plants suggest glutathione polysulfides as potential substrates (16), while another proposed glutathione-coordinated 2Fe-2S clusters (17). Multiple physiological substrates have also been proposed (18), with distinct Fe and S-containing and S-only containing intermediates both being exported by the transporter and being used for distinct processes: the Fe and S containing intermediate for cytosolic Fe-S protein maturation and a distinct sulfur containing intermediate being used for tRNA thiolation (19). While the exact nature of the physiological sulfur-containing substrate continues to be debated (20), the effect of ABCB7 depletion on the downstream cytosolic and nuclear Fe-S proteins is clear and conserved among commonly studied model organisms (7). However, despite its seeming universal importance, our understanding of how Fe-S cluster biosynthetic pathways work in the medically important apicomplexan parasites is incomplete.

Apicomplexa are obligate intracellular parasites which cause diseases of global importance, such as malaria and toxoplasmosis. *Toxoplasma gondii*, the causative agent of toxoplasmosis, can infect most warm-blooded animals and cause life-threatening diseases in the immunocompromised. As well as being an important pathogen, it is a versatile model organism for studying apicomplexan biology. Recent studies have begun to explore the understudied Fe-S biogenesis pathways in *Toxoplasma*. Studies looking at proteins in the CIA, ISC and SUF pathways have shown their importance for parasite fitness (21–23). Interruption of the ISC pathway, by depleting the mitochondrial scaffold protein ISU1, was shown to have a severe, but reversible, impact on parasite fitness and to trigger partial stage-conversion into bradyzoites, a cyst-enclosed persisting stage that is involved in the chronic phase of toxoplasmosis (21). The important relationship between the mitochondrion and cytosolic Fe-S biogenesis was further supported by a study which uncovered the unusual mitochondrial anchoring of the CIA scaffold protein NBP35, which was characterised as a cytosolic protein in other Eukaryotes (23). This unusual finding highlights how little we understand about how Fe-S cluster biogenesis intersects with parasite metabolism despite its importance to their fitness and life cycle.

Here, we wanted to understand the importance of the mitochondrial transporter ABCB7 for parasite fitness and to identify cytosolic Fe-S client proteins dependent on ABCB7 for activity.

## Results

### TGGT1_269000 encodes the *Toxoplasma* ABCB7

To identify the *Toxoplasma* homolog of the mitochondrial transporter ABCB7, we performed BLAST homology searches on the ToxoDB.org database (24) using both yeast and human homologs (ATM1, Uniprot: P40416; ABCB7, Uniprot: O75027). This approach identified TGGT1_269000 as the top hit (e-value: 9 x 10^-128^ percent identity: 35.16; and e-value: 1 x 10^-132^ percent identity: 35.17 respectively). While this gene had previously been identified as an ABCB7 homolog (21) it had also been suggested to be a putative homolog of the ABC transporter ABCB6 (25) which may have a role in porphyrin transport for haem biosynthesis. To investigate further, we performed phylogenetic analysis of TGGT1_269000 with ABCB6 and 7 from various species, which showed clear clustering of TGGT1_269000 with ABCB7 homologs (Fig. S1A). Multiple sequence alignment showed conservation of key functional residues involved in glutathione and ATP binding (Fig. S1B) and structural predictions using Alphafold (26, 27) suggested overall structural conservation with experimentally determined structures (Fig. S1C). These data are consistent with TGGT1_269000 encoding the *Toxoplasma* homolog of the mitochondrial transporter ABCB7.

To biochemically characterise the ABCB7 homolog, we first created a C-terminal triple hemagglutinin (HA) epitope tagged version of the endogenous protein using a CRISPR-mediated homologous recombination approach, outlined in Fig. 1B, in the TATiΔ*ku80* parental strain (28). A clonal line, named ABCB7-HA, was isolated from a positive pool by serial dilution and confirmed by PCR (Fig. 1C). Immunoblot analysis of ABCB7-HA showed a clear and specific signal, migrating between the 95 and 130 kDa molecular weight marker (Fig. 1D), consistent with the predicted size of 125 kDa. ABCB7 forms homodimers to create functional transporters (7). We therefore performed native-PAGE and immunoblot analysis of proteins extracted from ABCB7-HA parasites. Size estimation compared to membrane-bound bovine mitochondrial complexes, suggested migration at 287 KDa, a size consistent with migration of the homodimer (Fig. 1E). Longer exposures show the presence of a second band migrating at 111 kDa, possibly showing migration of monomeric ABCB7 that have not yet been incorporated in a homodimer (Fig. S2A). ABCB7-HA parasites treated with SDS, a stronger detergent which breaks weaker protein-protein interactions, showed a single band migrating at the size of a monomer (Fig. S2B). To provide extra support for homodimer formation we introduced a second, C-terminally Ty-epitope tagged, version of the protein and performed reciprocal co-immunoprecipitation (co-IP) experiments. Ty signal was detected in the bound fraction of the HA IP, and HA signal detected in the bound fraction of the Ty IP, while no signal from the unrelated proteins TOM40 and CDPK1 was found in either bound fraction (Fig. S2C) indicating interaction between the two copies of the protein, and consistent with ABCB7 homodimer formation.

Previous proteomic studies have suggested a mitochondrial localisation for ABCB7. The protein was detected in a recent *Toxoplasma* organelle proteomic atlas using hyperLOPIT and was predicted to be in the mitochondrial membranes fraction (29). It was also detected in a proximity-labelling proteome of the mitochondrial matrix (30) and in a complexome profiling analysis of *Toxoplasma* mitochondria (31). To confirm that ABCB7 is mitochondrially localised we performed an immunofluorescence assay (IFA). As predicted, the HA signal co-localised with the mitochondrial marker protein MYS (32), confirming mitochondrial localisation (Fig. 1F).

The ABCB7 sequence is predicted to have multiple transmembrane domains, using the prediction software TMHMM 2.0 (33), consistent with its function as a transporter. To provide support for ABCB7 being a transmembrane protein we performed sodium carbonate (Na_2_CO_3_) extractions. Resistance to Na_2_CO_3_ extraction can distinguish integral membrane proteins from soluble or periphery-associated membrane proteins (23, 34, 35). ABCB7-HA was detected in the Na_2_CO_3_ extraction-resistant pellet fraction, along with the mitochondrial integral membrane protein TOM40 (Fig. 1G), whilst the cytosolic protein CDPK1 and the peripherally membrane-associated ATP synthase subunit, ATPβ were soluble upon Na_2_CO_3_ treatment. To test that none of the proteins detected are inherently insoluble, samples were also subjected to extraction with the Triton X-100 (TX-100) detergent: as expected, in these conditions all proteins were found in the soluble fraction, indicating membrane proteins were efficiently solubilised in protein-detergent mixed micelles (Fig. 1G). Taken together, these data are in support of ABCB7 being an integral membrane protein, consistent with its role as a transporter.

Overall, we assigned TGGT1_269000 as the *Toxoplasma* homolog of the transporter ABCB7, and showed it is an integral membrane protein, likely forming homodimers, and localising to the mitochondrion.

### ABCB7 is required for growth and its depletion leads to partial conversion into bradyzoites

We next wanted to study the role of ABCB7 in *Toxoplasma* growth and Fe-S cluster metabolism. Data from a previous *Toxoplasma* genome-wide CRISPR screen (36), assigned ABCB7 a phenotype score of –4.96, indicating it is likely essential for the fitness of tachyzoites (the fast-replicating stage of the parasite responsible for the acute phase of toxoplasmosis) under standard culture conditions. Therefore, to study the role of this protein we generated a conditional knockdown line by replacing the native promoter with an anhydrotetracycline (ATc) regulated promoter (28, 37) using a CRISPR-mediated homologous recombination approach, in the ABCB7-HA background, outlined in Fig. 2A. A clonal line, named cKD-ABCB7-HA, was isolated from a positive pool by serial dilution and confirmed by PCR (Fig. 2B). To assess the extent of downregulation of ABCB7 transcript upon addition of ATc, we performed qRT-PCR on parasites grown in the presence or absence of ATc for two days. Parasites grown in ATc showed significantly lower levels of transcript, confirming controlled depletion (Fig. 2C). To assess the effect of depletion of the transcript on ABCB7 protein levels we performed immunoblot analysis on parasites grown in the absence or presence of ATc for one, two and three days. The ABCB7-HA protein was decreased after one day in ATc, and virtually undetectable after two and three days (Fig. 2D,E). Similar results were seen with IFA analysis (Fig. 2F). Notably, the levels of ABCB7 protein were ∼four times increased in the conditional knockdown line compared to the parental ABCB7-HA line, likely due to mis-regulation by inserting a non-native promoter (Fig. 2D,E). However, quantification of immuno-signal shows that after addition of ATc, protein levels are decreased compared to the parental cell line (Fig. 2E).

**Fig 2.**
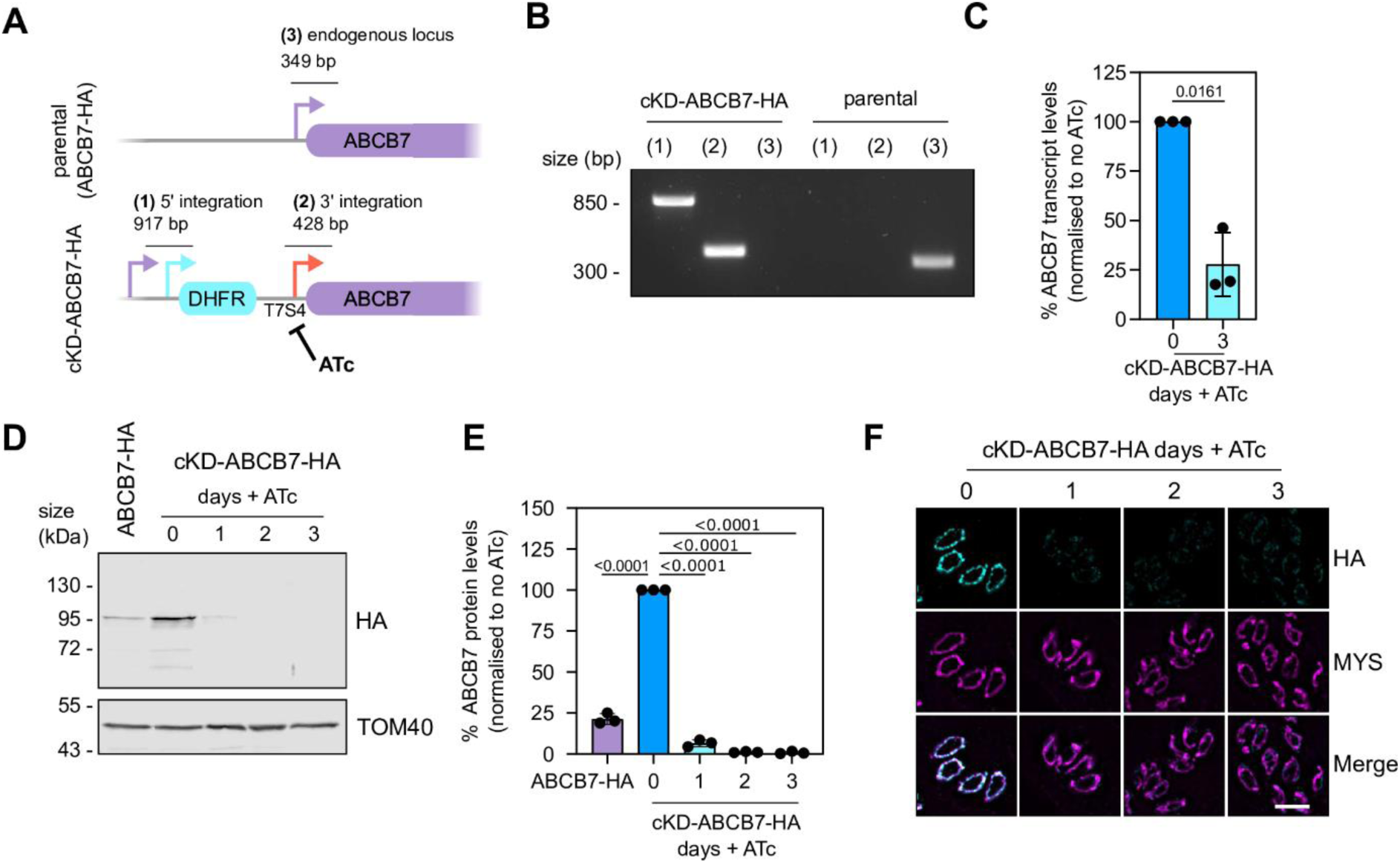
Replacement of *ABCB7* promoter with a regulatable promoter results in a tightly controlled conditional knockdown line. (A) Schematic of the promoter replacement strategy used to create a conditional knockdown of ABCB7 (cKD-ABCB7). The expected size of integration PCRs are shown. (B) PCR to test integration of DHFR selection cassette and the regulatable promoter into the endogenous locus, as outlined in *(A)*. (C) Relative transcript levels of ABCB7 in cKD-ABCB7-HA line after 3 days Anhydrotetracycline (ATc) treatment, measured by RT-qPCR. Error bars are mean -/+ S.D., and a one-sample t-test used to compare transcript levels in plus ATc to no ATc, which was set to one, n=3. (D) Immunoblot analysis of whole cell lysate extracted from cKD-ABCB7-HA parasites treated with ATc for zero, one, two and three days, and ABCB7-HA parasites. Samples were separated by SDS-PAGE, blotted, and detected using anti-HA, to visualise ABCB7-HA, and anti-TOM40 as a loading control. (E) Quantification of immunoblots in *(D)*. Each point represents a replicate, normalised to TOM40, and 0-day ATc set at 100%. Error bars are mean -/+ S.D., protein levels were compared to cKD-ABCB7-HA without ATc using one-way ANOVA, with a Dunnett correction for multiple comparisons, n = 3. (F) Immunofluorescence assay analysis of cKD-ABCB7-HA parasites treated with ATc for zero, one, two and three days, labelled with anti-HA to detect ABCB7-HA protein (cyan), and MYS as a mitochondrial marker (magenta). Scale bar is 5 µM.

The ABCB7 transporter is functionally required for the Fe-S cluster biosynthesis pathways in other species (4) and, combined with the low phenotype score (36), ABCB7 is predicted to be important for *Toxoplasma* fitness. To test this, we performed plaque assays. The cKD-ABCB7-HA line was grown on an HFF monolayer for eight days and were unable to form plaques in the presence of ATc (Fig. 3A), confirming the importance of ABCB7 for parasite growth *in vitro*. The cKD-ABCB7-HA parasites formed slightly smaller plaques than the parental control (Fig. 3A,B), suggesting a small fitness consequence of the insertion of the regulatable promoter, possibly due to increased ABCB7 protein levels (Fig. 2D,E). To show that the observed phenotype is due to the depletion of ABCB7 protein, we engineered a line containing a second Ty epitope-tagged copy of ABCB7 under the TUB8 promoter (38), in the conditional knockdown line, which we named cKD-ABCB7-HA/ ABCB7-Ty. IFA analysis showed this second copy is mitochondrially localised (Fig. S3A). Immunoblot analysis shows that after growing cKD-ABCB7-HA/ ABCB7-Ty in the presence of ATc for three days, the ATc regulatable HA-tagged version of the protein is depleted, but the Ty-epitope tagged version is unaffected (Fig. S3B). We performed plaque assays with the cKD-ABCB7-HA/ ABCB7-Ty line that showed no growth defect after eight days in ATc, demonstrating phenotypic rescue (Fig. 3A,B).

**Fig 3.**
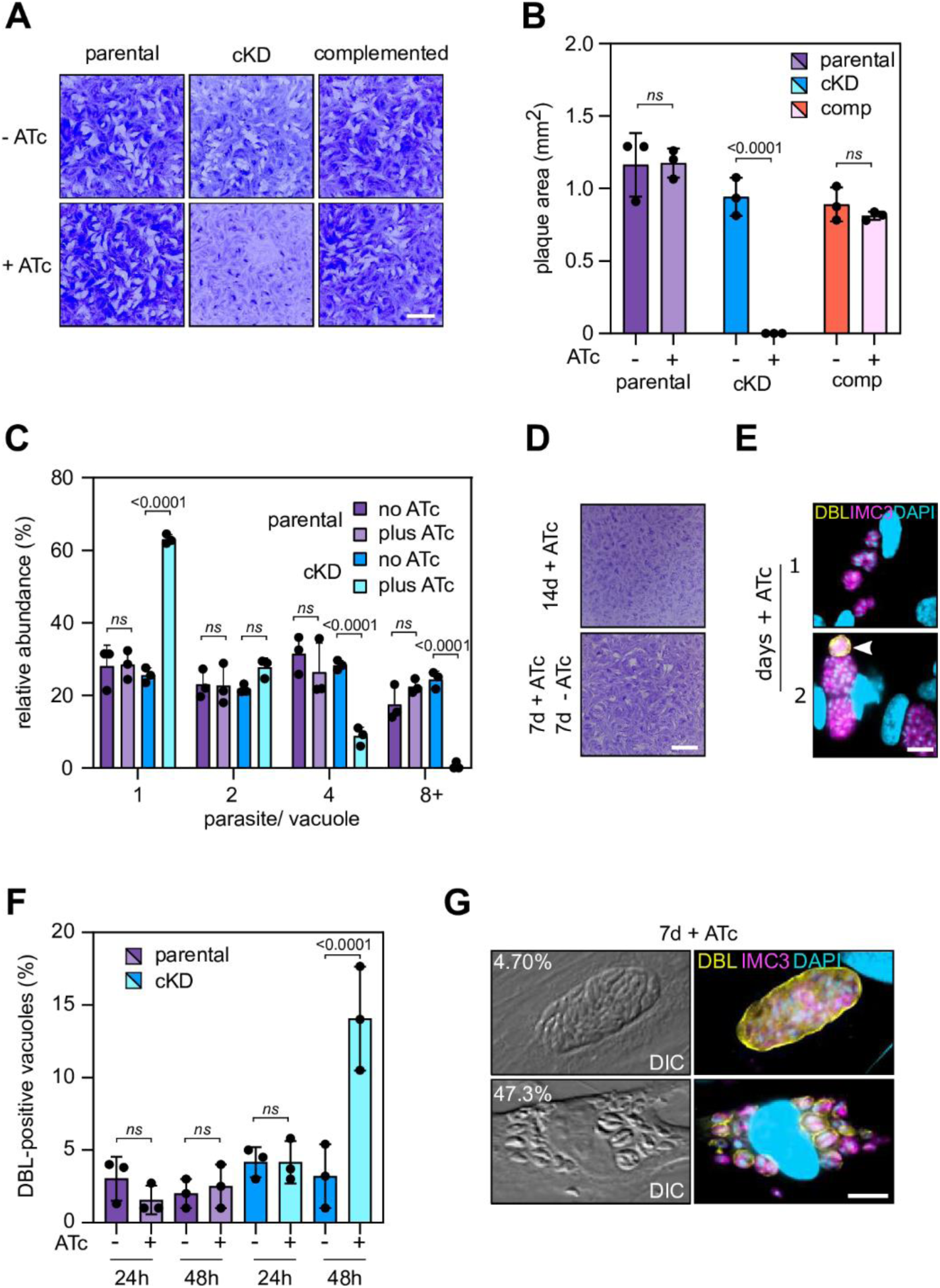
Downregulation of ABCB7 leads to a decrease in parasite replication and triggers partial parasite differentiation. (A) Plaque assays of parental, cKD-ABCB7-HA, and complemented parasites, grown in the presence or absence of ATc for 8 days. Scale bar is 5 mm (B) Quantification of plaque assays from *(A)*. Three independent experiments were performed, and 50 plaques measured per replicate. Mean of each replicate is displayed, ± SD. One way ANOVA followed by Turkey’s multiple pairwise comparisons was performed, and p-value from relevant pairs displayed. (C) Quantification of number of parasites per vacuole for parental and cKD-ABCB7-HA parasites. Parasites were grown in the presence or absence of ATc for 2 days before inoculation into fresh HFFs cells and grown for a further day in the presence or absence of ATc. Error bars are mean -/+ S.D from three independent experiments, for which over 100 vacuoles were counted for each replicate. One way ANOVA followed by Turkey’s multiple pairwise comparisons was performed, and p-value from relevant pairs displayed. (D) Plaque assays of cKD-ABCB7-HA parasites, grown for either fourteen days in the presence of ATc, or seven days in ATc followed by an ATc washout, followed by growth for seven days in the absence of ATc. Scale bar is 5 mm (E) Immunofluorescence assay of cKD-ABCB7-HA parasites treated with ATc for one or two days, labelled with IMC3 (magenta) to detect parasites and a lectin of *Dolicos biflorus* (DBL) to outline nascent cyst walls. 4′,6-diamidino-2-phenylindole (DAPI) was used to stain DNA. Scale bar is 10 µM. (F) Quantification of DBL-positive vacuoles from parental and cKD-ABCB7-HA parasites treated with ATc for one or two days. Data are from n = 3 independent experiments. One hundred vacuoles counted per experiment. Values are mean -/+ S.D. One way ANOVA followed by Turkey’s multiple pairwise comparisons was performed, and p-value from relevant pairs displayed. (G) Immunofluorescence assay of cKD-ABCB7-HA parasites treated with ATc for seven days, labelled with anti-IMC3 (magenta) to detect parasites and DBL to outline nascent cyst walls (yellow). Percentage at the top of the differential interference contrast (DIC) image represents the proportion of vacuoles that are DBL-stained and resemble mature cysts (top) or are DBL-stained but with four than four parasites (bottom), data are from n = 3 independent experiments in which at least 100 vacuoles were counted. DAPI was used to stain DNA. Scale bar is 10 µM.

To further characterise the effect of ABCB7 depletion on parasite growth, we grew parental and cKD-ABCB7-HA parasite in the presence or absence of ATc for two days, then allowed parasites to invade a HFF monolayer, grow for a further 24 hours, and then performed an IFA to assess parasite replication.

We quantified the number or parasites per vacuole and found a significant decrease in the number of vacuoles containing 4 and 8 (or more) parasites, and an increase in one parasite vacuoles, in the cKD-ABCB7-HA cell line grown in ATc (Fig. 3C). All other conditions resulted in similar numbers of parasites per vacuole. These data further support a marked impact on parasite replication when ABCB7 is depleted.

Our previous study looking at the mitochondrial Fe-S scaffold ISU1 (21) discovered a reversible growth defect when ISU1 is depleted. To assess if this was also the case in our ABCB7 depletion mutant, we grew cKD-ABCB7-HA parasites in ATc for fourteen days or ATc for seven days, followed by ATc wash-out and a further seven days in normal media. The parasites grown in ATc for fourteen days formed no plaques, whereas the parasites where the ATc was washed out formed small plaques (Fig. 3D). This suggests that, similar to what was observed for the ISU1 mutant, cKD-ABCB7-HA parasites are still viable after seven days of protein depletion and the growth defect is partially reversible. This phenotypic overlap with the ISC pathway mutant supports a connected role for these two pathways.

We have previously shown that disruption of mitochondrial functions, including mitochondrial Fe-S biosynthesis, led to partial differentiation into bradyzoites and started forming cyst-like structures (21, 35). To see if this was also the case for the ABCB7 conditional knockdown line we performed an IFA, using a lectin from *Dolichos biflorus* (DBL), which labels the cyst-wall glycoprotein CST1 that is built up during tachyzoite to bradyzoite conversion (39). We observed numerous DBL-positive structures (Fig. 3E), and quantification showed they were significantly increased in cKD-ABCB7-HA parasites grown in ATc for two days, with ∼15% of DBL positive vacuoles (Fig. 3 F). Long-term *in vitro* cultured cysts are usually large structures containing dozens of bradyzoites. To see whether ABCB7 depletion led to mature cysts, we performed DBL imaging after 7 days of incubation with ATc. More than half (52%) of the vacuoles showed DBL-staining, yet the vast majority of these (∼47%) contained four or fewer parasites, whilst only ∼5% formed large-disorganised cyst-like structures (Fig. 3G). These data suggest that, similar to what is seen in mitochondrial mutants, depletion of ABCB7 leads to the initiation of stage conversion, but this process is likely not completed.

Overall, our data show that ABCB7 is essential for parasite growth *in vitro*, and that its depletion leads to the initiation of partial stage conversion.

### Mitochondrial functions are unaffected after ABCB7 downregulation

Numerous studies in other model systems show that mutants in ABCB7 function leads to defects in cytosolic and nuclear Fe-S proteins, whilst mitochondrial Fe-S proteins remain unaffected. If this is the case in *Toxoplasma* too, we would expect mitochondrial biology to remain unaffected. To address this hypothesis, we first analysed general mitochondrial morphology upon depletion of ABCB7. Typically, intracellular parasites have a single mitochondrion which is predominantly found in a “lasso” shape that spans the parasite periphery. Our previous studies have shown that some defects in mitochondrial biology can result in alteration of this morphology (37, 40). We performed IFAs on parasites grown in the absence or presence of ATc for three days from parental and cKD-ABCB7-HA lines, as well as cKD-VDAC-HA, a conditional knockdown line of the mitochondrial transporter VDAC, which has a severe defect in mitochondrial morphology (40) and thus used as a positive control. This analysis showed in cKD-ABCB7-HA parasites, normal “lasso” shaped mitochondria can still be formed after three days in ATc, in contrast to cKD-VDAC-HA which has almost exclusively abnormal (i.e. shortened or collapsed) mitochondria (Fig. 4 A, B).

**Fig 4.**
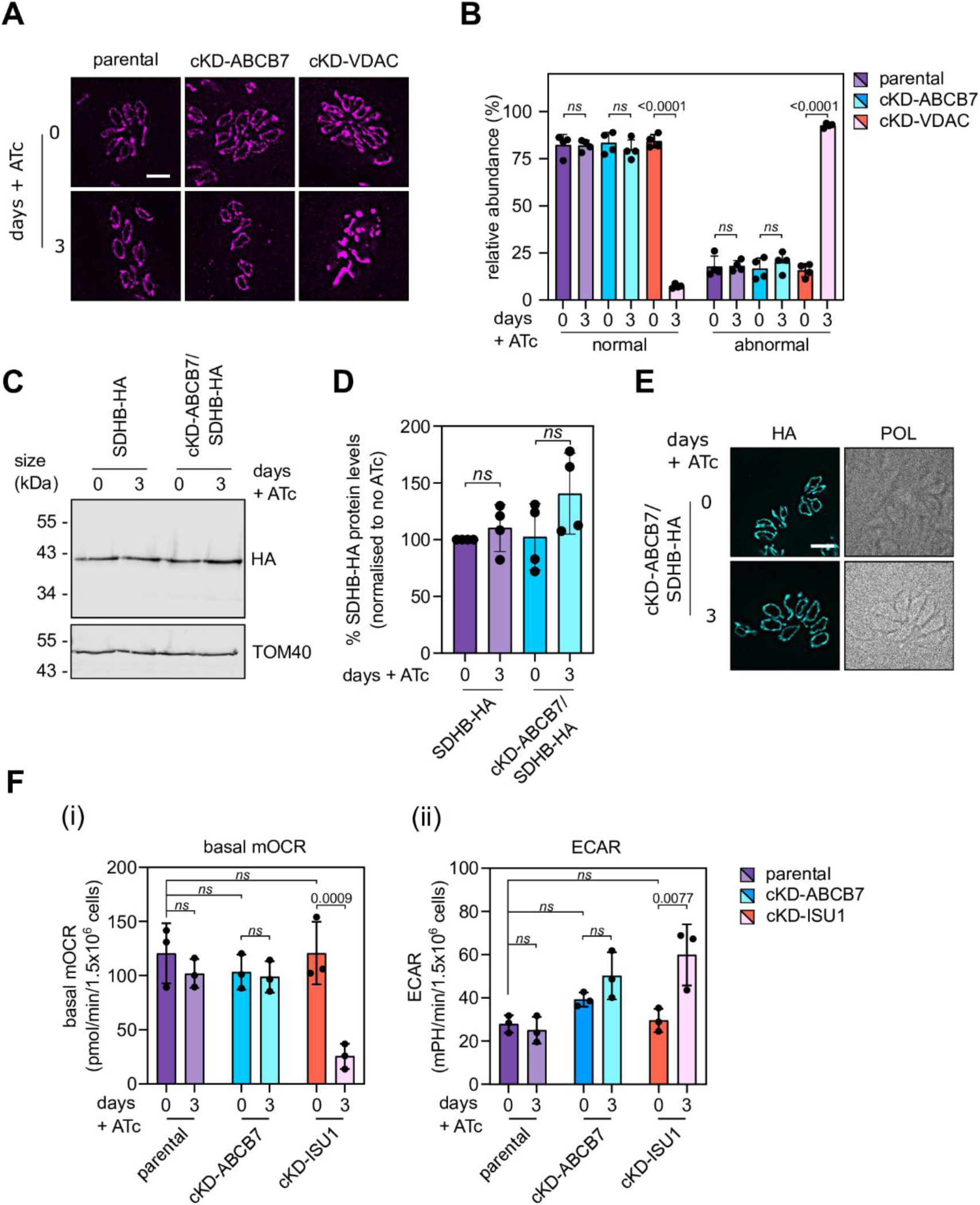
Mitochondrial functions are unaffected after downregulation of ABCB7. (A) Immunofluorescence assay analysis of parental, cKD-ABCB7-HA and cKD-VDAC-HA parasites grown in the presence or absence of ATc for 3 days, labelled with TOM40 to visualise mitochondrial morphology. (B) Quantification of mitochondrial morphology from parental, cKD-ABCB7-HA and cKD-VDAC lines at 3 days in the presence or absence of ATc. Morphologies were scored as normal or abnormal. Error bars are mean -/+ S.D from 4 independent experiments, for which over 150 vacuoles were counted for each replicate. One way ANOVA followed by Turkey’s multiple pairwise comparisons was performed, and p-value from relevant pairs displayed. (C) Immunoblot analysis of whole cell lysate extracted from SDHB-HA and cKD-ABCB7/SDHB-HA parasites grown in the presence or absence of ATc for 3 days. Samples were separated by SDS-PAGE, blotted, and detected using anti-HA, to visualise SDHB-HA, and anti-TOM40 as a loading control. (D) Quantification of immunoblots in *(C)*. Each point represents a replicate, normalised to TOM40, and SDHB-HA 0-day ATc was set at 100%. Error bars are mean -/+ S.D. from four independent experiments. One way ANOVA followed by Turkey’s multiple pairwise comparisons was performed, and p-value from relevant pairs displayed., n=4. (E) Immunofluorescence assay analysis of SDHB-HA and cKD-ABCB7/ SDHB-HA parasites grown in the presence or absence of ATc for three days, labelled with anti-HA to detect SDHB-HA protein. Scale bar is 5 µM. (F) Extracellular flux analysis, using a seahorse analyser, basal mitochondrial oxygen consumption rate (OCR) and extracellular acidification rate (ECAR) of parental, cKD-ABCB7-HA and cKD-ISU1-HA parasites grown in the presence or absence of ATc for three days. Graphs show mean -/+ S.D. from three independent experiments. One-way ANOVA followed by Turkey’s multiple pairwise comparisons was performed, and p-value from relevant pairs displayed.

Previous studies have shown that defects in the mitochondrial Fe-S biosynthetic pathway (the ISC pathway), either through depletion of the cysteine desulfurase NFS1 (23) or the scaffold protein ISU1 (21), leads to a decrease in the abundance of the Fe-S cluster containing succinate dehydrogenase (mETC complex II) subunit SDHB. While on the other hand, we expect that ABCB7 depletion which does not affect mitochondrial ISC in other systems, would likely be unaffected here too. To test this hypothesis, we created again a conditional knockdown line of ABCB7, as before, but now in a background where SDHB is C-terminally HA epitope tagged (31) (Fig. S4A). Immunoblot analysis shows that depletion of ABCB7 does not result in a decrease in SDHB protein (Fig. 4C,D). IFA analysis also shows no decrease or change in localisation of SDHB protein levels (Fig. 4E). Together these data show that depletion of ABCB7 does not decrease SDHB protein abundance, in sharp contrast to the decrease observed upon depletion of ISC pathway members ISU1 or NFS1.

One of the main clients of mitochondrially derived Fe-S clusters is the mETC, where complexes II, III and IV in *Toxoplasma* all contain subunits predicted to contain clusters (21, 30, 31, 41, 42). Depletion of NFS1 or ISU1 lead to a decrease in mETC capacity (21, 23). To test if depletion of ABCB7 affects the mETC, we measured the basal mitochondrial oxygen consumption rate (basal mOCR), using a seahorse extracellular flux analyser. The basal mOCR has previously been used in *Toxoplasma* as a proxy measurement for mETC activity (30, 35, 41, 43), and mutants in the mitochondrial Fe-S pathway have been shown to have a lower mOCR (23). We observed no decrease in basal mOCR in the cKD-ABCB7-HA line grown in ATc for three days (Fig. 4G (i)), again suggesting no defect in the mETC. The parental line was also unaffected upon ATc treatment, but growth in ATc resulted a significant decrease in basal mOCR in the cKD-ISU1-HA line as expected. We also measured the extracellular acidification rate (ECAR), as a proxy for general parasite metabolism. We observed no defect in ECAR, showing that the parasites are still viable (Fig. 4G (ii)). ECAR in cKD-ISU1-HA was increased, perhaps due to a compensatory increase in glycolysis.

These data suggest that depletion of ABCB7 does not affect the important mitochondrial functions that Fe-S proteins are involved in, and thus that likely the biosynthesis of mitochondrial Fe-S proteins is unaffected. This suggests that the observed growth and differentiation defects of the cKD-ABCB7-HA mutant likely involve proteins or pathways that act downstream of the mitochondrion.

### Label free quantitative proteomics identifies pathways affected by ABCB7 depletion

Having shown the importance of ABCB7 for *Toxoplasma* fitness, we wanted to explore further the effect on ABCB7 depletion of the biosynthesis of Fe-S clusters, and on parasite metabolism. Our previous study on the importance of the ISC and SUF pathway on parasite fitness used label-free quantitative (LFQ) proteomics to assess the effect of disrupting specific proteins in the pathways on the global parasite proteome (21). We performed a similar analysis on parental and cKD-ABCB7-HA parasites grown in ATc for three days. Total protein samples from four independent experiments were subjected to mass spectrometry analysis, and LFQ values were calculated to compare protein levels between parental and knockdown parasites. 3922 proteins were detected at high enough quality to calculate LFQ intensity values. The ABCB7 protein was detected in all parental replicates, but were undetected in all mutant replicates, confirming it is efficiently downregulated below the detection threshold. 140 proteins were significantly increased in abundance (log2(fold change) ≥ 0.55) in the ckD-ABCB7-HA line compared to parental, whilst 249 were of lower abundance (log2(fold change) ≤ - 0.55) (Table S1,2). There was little overlap in the identity of the proteins increased or decreased in abundance compared to similar analysis on the mitochondrial Fe-S scaffold ISU1 (Fig. 5B) or the apicoplast NFS2 (Fig. 5C) (21), suggesting the pattern of protein abundance change is specific to the depletion of ABCB7. Analysis of the predicted localisation of decreased proteins through the hyperLOPIT-based organelle proteomic dataset shows a large number that are in the cytosolic or nuclear compartments (158 out of 256) (Fig. 5D), consistent with what might be expected if ABCB7 function is important for the cytosolic FeS cluster assembly pathway. In addition, increases in protein abundance were found in two other compartments, the dense granule (38 of 140) and mitochondria (27 of 140) (Fig. 5E).

**Fig 5.**
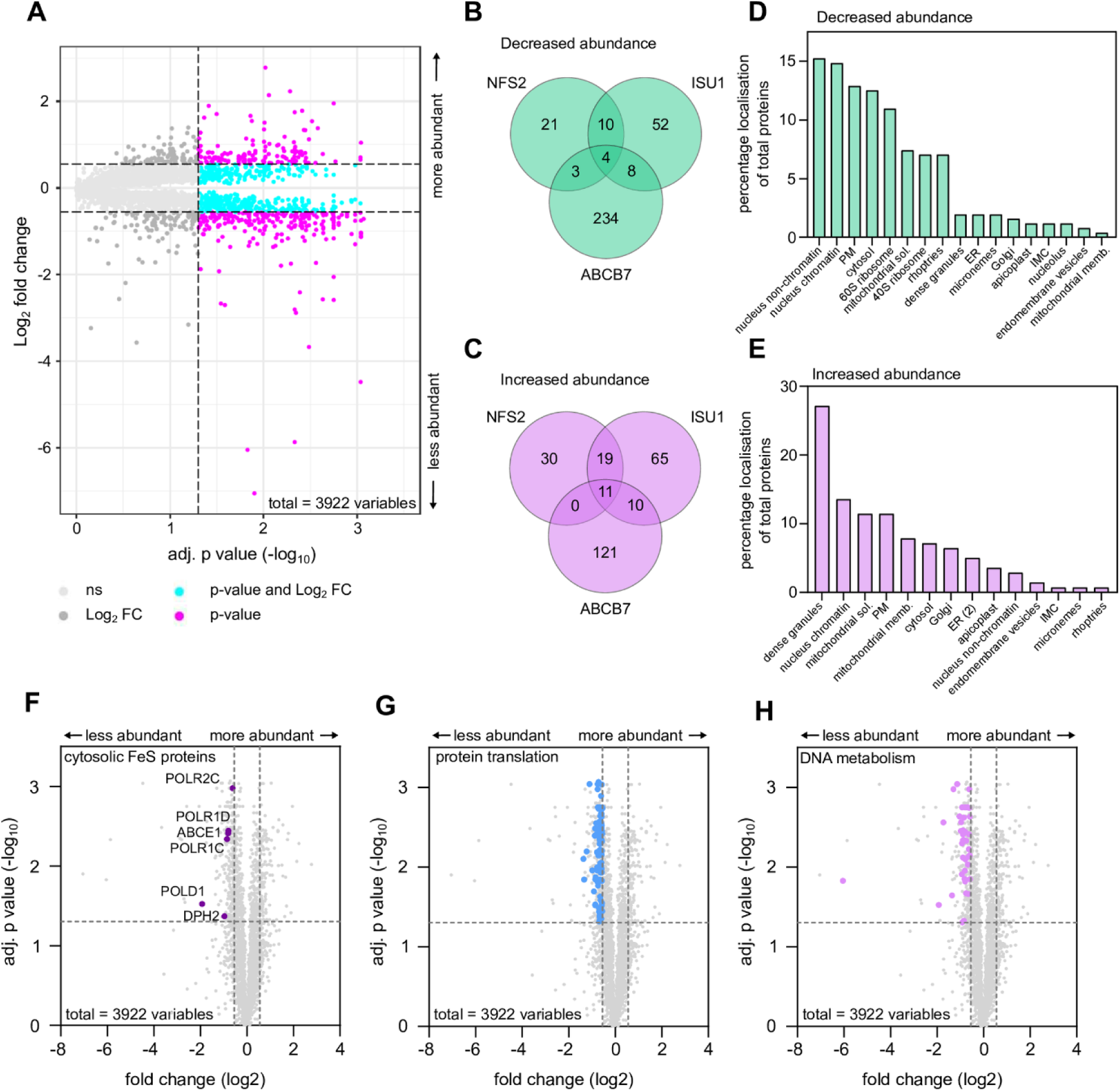
ABCB7 depletion impacts cytosolic Fe-S proteins and proteins involved in DNA metabolism and translation. (A) Volcano plot showing the difference in protein abundance, determined by label-free quantitative proteomic data, in cKD-ABCB7-HA parasites grown in ATc for three days. Y-axis shows the Log_2_ fold change compared to the parental line grown in the same conditions, and the X axis shows the -log10(p value) after a student’s t-test with a Benjamini correction for multiple comparisons applied, for four independent biological replicates. (B) and (C) Venn diagram depicting shared and unique proteins whose abundance is affected by the depletion of ABCB7, and NFS2 and ISU1 from (21). (D) and (E) Localisation, predicted by HyperLOPIT (29), of proteins that are decreased (*D*) or increased (*E*) upon depletion of ABCB7. (F) Volcano plot of LFQ data with cytosolic Fe-S proteins showing decreased abundance labelled. (G, H) Volcano plot of LFQ data with proteins involved in protein translation (G) and DNA metabolism (H) showing decreased abundance labelled.

To investigate in more detail the effect of depletion of ABCB7, we looked in our proteomics dataset for components of the ISC and CIA pathway. We did not observe a strong effect on the proteins of the CIA pathways, with none of its members increased or decreased in abundance above the ≥ 0.55 threshold (Fig. S5A). This suggests that ABCB7 depletion has little impact on the CIA pathway abundance.

Interestingly, in the mitochondrial ISC pathway, two proteins involved in early steps of sulfur generation, the cysteine desulfurase NFS1 and ISD11, were increased in abundance, as well as a glutaredoxin (Fig. S5A), perhaps upregulated as part of a feedback loop to compensate for the lack of sulfur-containing intermediate in the cytosol.

We previously predicted non-organellar Fe-S proteins using the Fe-S protein prediction software MetalPredator (21, 44). We manually updated this to include proteins predicted to be organellar in the hyperLOPIT dataset (29), but which have experimental evidence to be cytosolic facing proteins, for example NBP35 (23), RlmN (45) and ELP3 (46). In the absence of a functional cluster, Fe-S cluster containing proteins may be degraded (47). 18 of the 28 proteins in this list were detected at high enough quality to calculate LFQ intensity values. Six of these were of decreased abundance, below the fold change cut-off (Table 1; Fig. 5F, S5B). These include Fe-S cluster containing subunits of DNA and RNA polymerases, as well as factors important for translation. In addition, the predicted Fe-S cluster containing DNA primase large subunit, TGGT1_297840 (PRIM2) and the DNA polymerase family B protein TGGT1_319860 (POLE), were detected in all four parental replicates, but not in all replicates of the ABCB7 knockdown sample (Table 1; Table S1,2). In contrast, none of the predicted mitochondrial or apicoplast Fe-S proteins were found to be decreased upon ABCB7 depletion (Fig. S5C,D). These data suggest that a depletion of ABCB7 leads to a preferential decrease in abundance of non-organellar Fe-S proteins, as expected when the cytosolic biosynthetic pathway is disrupted.

**Table 1:** Cytosolic and nuclear Fe-S proteins decreased in abundance upon ABCB7 depletion.

| Gene ID | Name | Phenotype score | Localisation (hyperLOPIT) | Function |
| --- | --- | --- | --- | --- |
| TGGT1_258030 | POLD1 | -4.45 | nucleolus | Subunit of DNA polymerase delta complex with role in DNA replication |
| TGGT1_238190 | POLR1C | -4.42 | nucleus - non-chromatin | Subunit of DNA-directed RNA polymerase I and III |
| TGGT1_261540 | POLR1D | -3.54 | nucleus - non-chromatin | Subunit of DNA-directed RNA polymerase I and III |
| TGGT1_267390 | POLR1C | -5.2 | nucleus - non-chromatin | Subunit of DNA-directed RNA polymerase I and III |
| TGGT1_216790 | ABCE1 | -4.11 | nucleus - non-chromatin | Ribosome recycling factor |
| TGGT1_261060 | DPH2 | -1.94 | n.d. | Enzyme required for diphthamide biosynthesis; a post-translational modification important for translational fidelity |
| TGGT1_297840 | PRIM2 | -4.86 | n.d. | Subunit of DNA primase and DNA polymerase alpha complex with role in DNA synthesis |
| TGGT1_319860 | POLE1 | -3.85 | n.d. | Subunit of DNA polymerase epsilon complex with role in DNA replication |

To look at the effect of ABCB7 depletion on parasite biology we further inspected the LFQ proteomic data, looking for pathways that were decreased in abundance. A gene ontology enrichment analysis (ToxoDB) of the 249 proteins decreased in abundance indicated that proteins involved in translation (59 proteins, 6.31-fold enrichment, p-value 2.5 x 10^-33^), ribosome biogenesis (10 proteins, 3.93-fold enrichment, p-value 1.78 X 10^-4^), DNA replication (9 proteins, 4.39-fold enrichment, p-value 1.59 x 10^-4^), DNA replication initiation (6 proteins, 16.26-fold enrichment, p-value 3.34 x 10^-7^) and DNA dependent DNA replication (6 proteins, 8.61-fold enrichment, p-value 3.73 x 10^-5^) were significantly enriched in our dataset. Further investigation of poorly annotated proteins in our dataset using HHpred and domain searches found more factors involved in protein translation, ribosome and tRNA biogenesis, DNA and RNA replication and metabolism. In total, we found 133 proteins that were significantly decreased below the fold change cut-off with homology to known components of these processes (Fig. 5G,H, Table S1,2) consistent with experimental observations from other organisms (48).

### ABCB7 depletion leads to a decrease of the DNA polymerase subunit POLD1 and the ribosome biogenesis factor ABCE1

To validate our proteomic data, we decided to perform focused experiments on two Fe-S cluster containing proteins that were significantly decreased. One of the most decreased proteins was the Fe-S cluster containing DNA polymerase delta catalytic subunit POLD1 (TGGT1_258030). POLD1 in humans has previously been shown to interact with the CIA-targeting complex and depend on MMS19 for cluster insertion (48). To study the effect of ABCB7 depletion on POLD1, we created a C-terminal triple HA epitope tagged version of the endogenous protein, using the same strategy as previously described and as outlined in Fig. S4B and named the line POLD1-HA. Immunoblot analysis of POLD1-HA showed a clear and specific band (Fig. 6A), migrating between the 130 and 170 kDa molecular weight marker, consistent with the predicted size of 143 kDa. IFA showed a diffuse signal across the cell, with an area of high signal overlapping with DAPI (Fig. 6B). Next, we created a conditional knockdown line of ABCB7 in the POLD1-HA background (Fig. S4D), named cKD-ABCB7/ POLD1-HA. To test if ABCB7 depletion affects POLD1-HA protein levels we performed an IFA on cKD-ABCB7/ POLD1-HA, grown in the presence or absence of ATc for three days. HA signal was severely decreased, and largely absent from the nucleus, in the parasites grown in ATc (Fig. 6C). Immunoblot analysis also showed a decrease in POLD1 protein level in cKD-ABCB7/POLD1-HA parasites grown in ATc (Fig 6D,E). These data suggest that depletion of ABCB7 leads to a decrease in POLD1 protein levels, consistent with the requirement of ABCB7 for cytosolic Fe-S cluster maturation.

**Fig 6.**
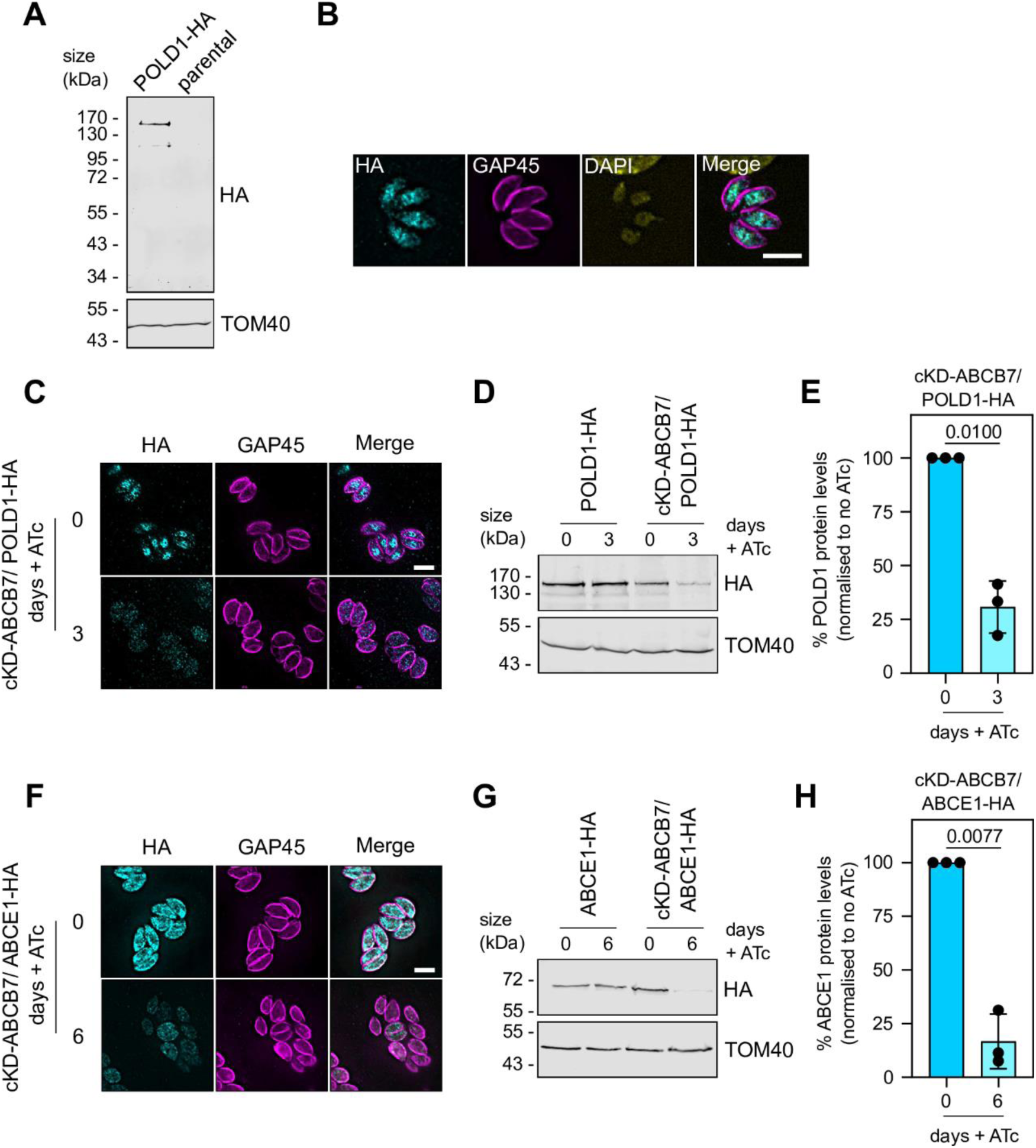
ABCB7 depletion impacts POLD1 and ABCE1 protein levels. (A) Immunoblot analysis of whole cell lysate extracted from POLD1-HA and parental parasites. Samples were separated by SDS-PAGE, blotted, and detected using anti-HA, to visualise POLD1-HA, and anti-TOM40 as a loading control. (B) Immunofluorescence assay analysis of POLD1-HA parasites, labelled with anti-HA to detect POLD1-HA (cyan), anti-GAP45 (magenta) to outline the parasites, and DAPI stan to label the nucleus. Scale bar is 5 µM. (C) Immunofluorescence assay analysis of cKD-ABCB7/ POLD1-HA parasites, grown in the presence or absence of ATc for three days, labelled with anti-HA to detect POLD1-HA (cyan), anti-GAP45 to outline the parasites (magenta). Scale bar is 5 µM. (D) Immunoblot analysis of whole cell lysate extracted from POLD1-HA and cKD-ABCB7/POLD1-HA parasites grown in the presence or absence of ATc for three days. Samples were separated by SDS-PAGE, blotted, and detected using anti-HA, to visualise POLD1-HA, and anti-TOM40 as a loading control. (E) Quantification of cKD-ABCB7/ POLD1-HA immunoblots in *(D)*. Each point represents a replicate, normalised to TOM40, and cKD-ABCB7/POLD1-HA zero-day ATc set at 100%. Error bars are mean -/+ S.D., and a one-sample t-test used to compare protein levels in plus ATc to no ATc, n = 3. (F) Immunofluorescence assay analysis of cKD-ABCB7/ ABCE1-HA parasites, grown in the presence or absence of ATc for six days, labelled with anti-HA to detect ABCE1-HA (cyan), anti-GAP45 to outline the parasites (magenta). Scale bar is 5 µM. (G) Immunoblot analysis of whole cell lysate extracted from ABCE1-HA and cKD-ABCB7/ ABCE1-HA parasites grown in the presence or absence of ATc for six days. Samples were separated by SDS-PAGE, blotted, and detected using anti-HA, to visualise ABCE1-HA, and anti-TOM40 as a loading control. Representative image of three independent experiments. (H) Quantification of cKD-ABCB7/ABCE1-HA immunoblots in *(G)*. Each point represents a replicate, normalised to TOM40, and cKD-ABCB7/ABCE1-HA 0 -day ATc set at 100%. Error bars are mean -/+ S.D., and a one-sample t-test used to compare protein levels in plus ATc to no ATc, n = 3.

We next performed similar experiments on the Fe-S cluster containing ribosome recycling factor ABCE1, which has previously been shown to decrease in abundance upon disruption of the CIA pathway (23) and was found to be decreased in our proteomic data. We created a ABCE1-HA line (Fig. S4E,F) and showed similar protein size by immunoblot and localisation by IFA as previously published (Fig. S4G,H) (23). We engineered a conditional knockdown line of ABCB7 in the ABCE1-HA background (Fig. S4I) To test if ABCB7 depletion affects ABCE1-HA protein levels we performed IFA and immunoblot analysis on cKD-ABCB7/ ABCE1-HA, grown in the presence or absence of ATc. After three days in ATc, a slight decrease in ABCE1-HA levels in some cells could be detected by IFA (Fig. S4J). No detectable difference could be observed by immunoblot (Fig. S4K). This may be due to increased sensitivity of our proteomics approach compared to immunoblot analysis. To investigate further we performed the same experiments at a later (6-day) time point of incubation with ATc, where we saw that ABCE1-HA signal was severely decreased (Fig. 6F,G, H). As a control, we performed the same experiment on cKD-ABCB7/ SDHB-HA parasites at 6-day ATc growth, and saw no decrease in SDHB-HA signal (S4L Fig), indicating the decrease is specific for ABCE1 and does not reflect a general proteolysis due to a loss of viability, and is consistent with the parasites remaining viable for an extended period upon ABCB7 depletion (Fig. 3C,H). Taken together, these experiments confirm POLD1 and ABCE1 decrease in abundance after ABCB7 depletion, providing validation of our LFQ proteomic dataset and providing support for ABCB7’s role in cytosolic Fe-S maturation.

### ABCB7 depletion results in a global decrease in cytosolic translation

Given the observed decrease of many proteins involved in translation in our LFQ data (Fig. 5G), we decided to investigate if the overall cytosolic translation rate was affected in the ABCB7 mutant using puromycin incorporation assays. Puromycin is an amino-nucleoside antibiotic which blocks protein translation by incorporating itself into the nascent polypeptide chain during translation and blocking further synthesis. By measuring the puromycin incorporation, through immunoblot assays, the rate of protein synthesis can be measured (49). We first exposed freshly lysed parasites with either 100 µg/ml puromycin for 15 minutes; 100 µg/ml cycloheximide (a translation inhibitor), followed by 15 minutes with puromycin; and no treatment. We then looked at these parasites by immunoblot analysis with antibodies against puromycin. Substantial labelling had occurred in the puromycin incubated parasites, while none was seen in the untreated control, showing the specificity of the assay (Fig. 7A). Parasites incubated with cycloheximide prior to puromycin labelling showed much decreased labelling, showing that the assay can detect changes in translation. Next, to assess the rate of protein translation upon ABCB7 depletion we grew parental and cKD-ABCB7-HA for three days in the presence or absence of ATc, followed by puromycin labelling and immunoblot analysis. As a control we included cKD-ISU1-HA parasites, which have a severe defect in the mitochondrial ISC pathway, but for which similar LFQ proteomic analysis suggest no decrease in translation related components (21). Parasites depleted in ABCB7 showed a ∼40% decrease in puromycin incorporation (Fig. 7B,C), while parental or cKD-ISU1-HA parasites grown in ATc showed no decrease, suggesting that translation is decreased only when the CIA pathway is disrupted. Depletion of ISU1 results in a growth phenotype of similar severity to ABCB7, thus the decrease in translation defect is not a general phenotype of disrupted Fe-S cluster assembly, but rather a specific defect of ABCB7 depletion. A similar decrease in puromycin incorporation was seen in cKD-ABCB7-HA, but not parental, cells grown for 3 days in ATc in an immunofluorescence-based version of the assay (Fig. 7D).

**Fig 7.**
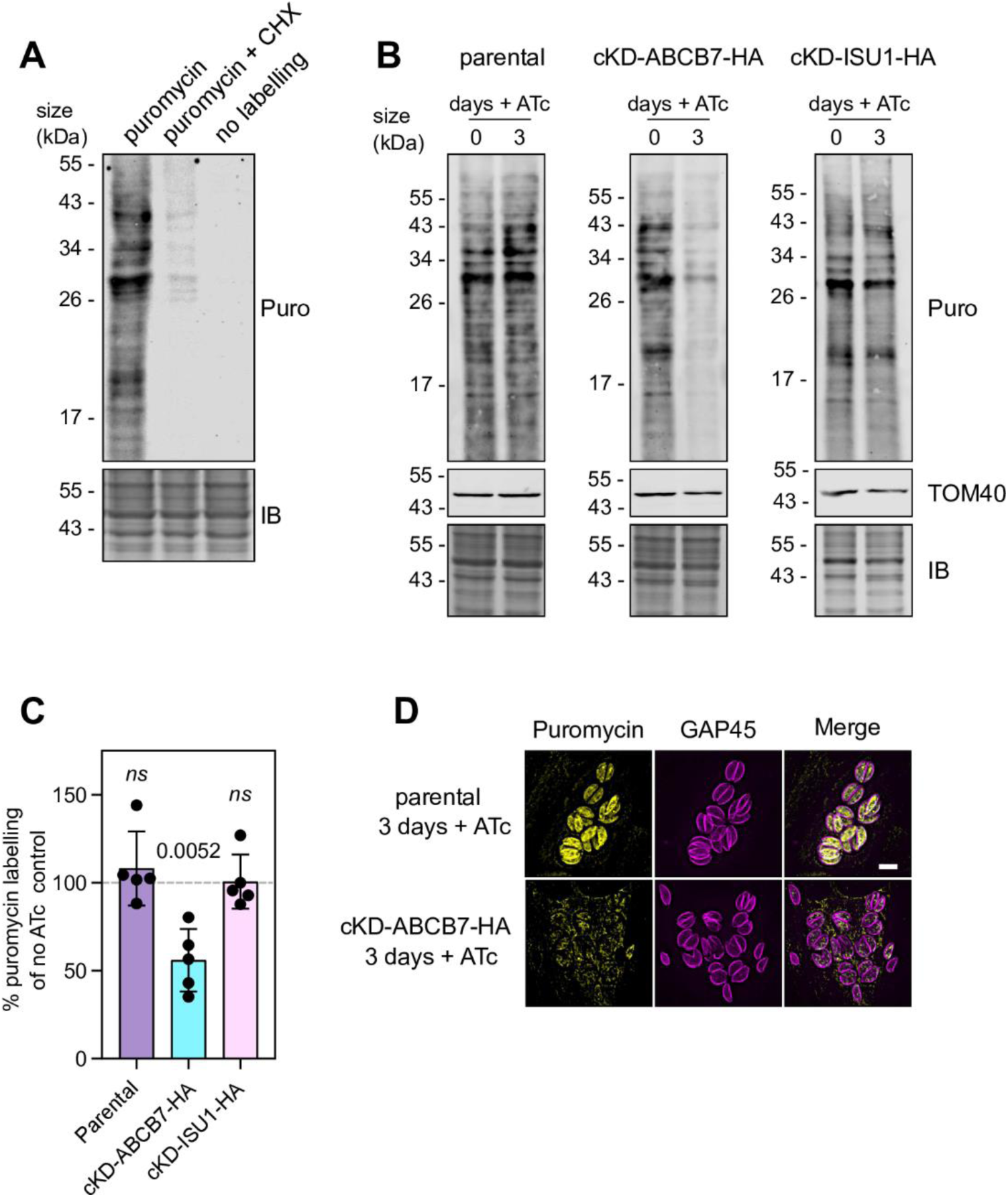
Cytosolic translation is decreased after depletion of ABCB7. (A) Immunoblot analysis of whole cell lysate incubated with (i) puromycin for 15 minutes, (ii) cycloheximide for 10 minutes, followed by puromycin for 15 minutes, and (iii) no treatments. Samples were separated by SDS-PAGE, blotted, and detected using anti-puromycin to detect puromycin incorporation (top panel), or stained with instant blue (bottom panel) as a loading control. (B) Immunoblot analysis of whole cell lysate from parental, cKD-ABCB7 and cKD-ISU1-HA, grown in the presence or absence of ATc for three days, and treated with puromycin for fifteen minutes. Samples were separated by SDS-PAGE and labelled as in *(B)*. (C) Quantification of immunoblots in *(C)*. Each point represents a replicate, normalised to instant blue signal, and no ATc set at 100%. Error bars are mean -/+ S.D., and a one-sample t-test used to compare levels in plus ATc to no ATc, n = 5. (D) Immunofluorescence assay analysis of parental and cKD-ABCB7-HA parasites, grown in the presence of ATc for three days, labelled with anti-puromycin (yellow), anti-GAP45 to outline the parasites (magenta). Scale bar is 5 µM.

The decrease in abundance of proteins involved in translation and decrease in puromycin incorporation upon ABCB7 depletion suggests that disruption of the CIA pathway results in a moderate defect in the translation rate in the cell, which may be due to a decrease in abundance of translational components (Fig 5G), including potentially Fe-S cluster containing components of the translation machinery, such as ABCE1 and DPH2 (Table 1).

## Discussion

Fe-S clusters are ubiquitous and essential inorganic cofactors for numerous essential life processes that have dedicated biosynthetic pathways. Despite some variation in different lineages, the core machinery of these biosynthetic pathways is remarkably conserved. Recently studies on Fe-S cluster biosynthesis in the important eukaryotic pathogen *Toxoplasma gondii*, have revealed the presence and functional importance of components from the ISC pathway in mitochondria (21, 23), the SUF pathway in the apicoplast (21, 22) and the cytosolic pathway (23). Understanding their role in *Toxoplasma*, and uncovering any lineage-specific features, is therefore important in the effort to combat these pathogens. These studies have also described some lineage-specific aspects of otherwise conserved components. For example, the cytosolic scaffold NBP35, which plays a central role in assembling cytosolic clusters, was found to be bound to the outer surface of the mitochondria (23), whereas in other model systems it is a soluble cytosolic protein, with one study suggesting a nuclear localisation (50). The exact reason for NBP35’s re-localisation remains unclear, but it may highlight the connection between the CIA and ISC pathways in apicomplexan parasites and reflects the complex evolutionary history of these proteins in eukaryotes (51–53)

Studies in a wide array of organisms clearly show that the role of the ABCB7 transporter in connecting the mitochondrial ISC and cytosolic CIA pathway is conserved (3). Downstream cytosolic Fe-S proteins are dependent on the cysteine desulfurase activity of the mitochondrial NFS1 enzymes, and the resulting sulfur intermediate that is transported out of the mitochondrial matrix by ABCB7. ABCB7 mutants in yeast and mammals are impaired in thio-modification of cytosolic tRNAs, protein translation, DNA replication and genome maintenance (3, 18, 48, 54, 55). This is consistent with what we find in our study, with LFQ proteomic data pointing to significant defects in translation and DNA metabolism, as well as Fe-S proteins involved in these processes (Fig. 5F-H; Table 1). Experiments also confirmed the specific depletion of the ribosome splitting factor ABCE1 and the DNA polymerase subunit POLD1. Further studies could test if thio-modification of tRNAs is affected. Growth assays (Fig. 3D) and ECAR measurements from extracellular flux analysis (Fig. 4F) show that parasites are still viable when these assays are performed and so act as a useful control that observed defects are specific and not due to a general decrease in parasite viability. Other studies of ABCB7 homologs, for example in plants, have assayed cytosolic Fe-S cluster defects through in-gel activity assays of enzymes containing Fe-S cluster such as aldehyde oxidase, xanthine dehydrogenase and cytosolic isoforms of aconitase (10, 56–58). *Toxoplasma* lacks obvious homologs of these enzymes or does not have clear cytosolic-only isoforms and so we were unable to perform similar analysis here.

We previously predicted *Toxoplasma gondii* to have 64 Fe-S cluster containing proteins (21) using the computational tool MetalPredator (44). However, predicting the ability of proteins to bind Fe-S clusters from sequence alone has long proved challenging (59) and other species contain many more Fe-S proteins. For example, a recent inventory of *E. coli* Fe-S proteins numbered 144 (59). In eukaryotes, up to 1% of the proteome is predicted to contain Fe-S clusters (60) (which would be 84 proteins if this trend is observed in *Toxoplasma*). Given these facts, it is plausible that *Toxoplasma* contains more, as yet unidentified, Fe-S proteins. While many Fe-S proteins are degraded if they do not bind to a cluster (47), not all proteins are destabilised. Our previous study using LFQ proteomics to study a mutant of the mitochondrial Fe-S cluster synthesis pathway highlighted a sharp decrease in abundance for Fe-S proteins of the mETC, SDHB, ApiCox13 and Rieske (21). While ABCB7 localises to the mitochondrion, LFQ analysis of the corresponding mutant highlighted a completely different profile, as the only MetalPredator-predicted candidates found to be less abundant were cytosolic or nuclear. This data both suggests that the changed proteins are bona fide Fe-S proteins, and confirms the involvement of *Toxoplasma* ABCB7 in the CIA pathway. Although LFQ is a useful approach for validation of clients that are highly responsive to lack of Fe-S cluster occupancy, as not all Fe-S proteins become unstable in the absence of the cofactor, more specialised techniques like chemoproteomics approaches will be required to uncover new Fe-S client proteins (59).

Our previous study on mitochondrial Fe-S biosynthesis found a major mutant phenotype to be the initiation of stage conversion into bradyzoites (21). This phenotype was also seen in other mitochondrial mutants, for example of the mETC or mitochondrial translation (21, 35). This phenotype was also seen in the ABCB7 mutant. The integration of environmental or metabolic cues that lead to conversion into bradyzoites is not fully understood. Stage conversion in the ABCB7 mutant may be triggered by metabolic signal derived from sensing a lack of functional Fe-S clusters in important enzymes. Another possibility, given the decreased abundance of many proteins involved in translation in the ABCB7 mutant, is that a block in translational could induce the parasite to begin differentiation. Previous studies have shown translational control to be an important aspect of initiating differentiation (61–65). More work needs to be done to understand how stress and metabolic deficiencies influence a parasite’s decision to initiate conversion.

ABCB7 is almost universally conserved in eukaryotes, and acts as a link between the ISC and CIA pathways in all studied examples. However, there are exceptions to be found in microbial parasites, especially those containing reduced mitochondria (mitochondria related organelles-MROs), which apparently lack ABCB7 homologs. The diplomonad intestinal parasite *Giardia*, which contains homologs of both CIA and ISC pathway components, lacks an ABCB7 homolog (66, 67), suggesting an alternative export route for the NFS1-generated sulfur intermediate (3), or raising the possibility that the two pathways are unconnected in this species The anaerobic protist *Pygsuia biforma*, lacks an ABCB7 homolog as well as ISC pathway components (68). Two amoebazoans, *Entamoeba histolytica* and *Mastigamoeba balamuthi*, lack ABCB7 homologs, as well as ISC components, instead utilising the bacterial-like NIF pathway (3). A more extreme example is the Oxymonad *Monocercomonoides exilis* which lacks a mitochondrion, and instead supplies its CIA pathway by a cytosolic SUF pathway (69, 70). These examples serve to highlight the biochemical diversity of non-canonical Fe-S assembly pathways present in the microbial parasites and highlight the utility of studying these pathways to gain a full picture Fe-S assembly in eukaryotes. While our data clearly demonstrate a conserved, canonical, function of ABCB7 in cytosolic Fe-S assembly, *Toxoplasma* may also possess unusual features in its Fe-S assembly pathways that remain to be discovered and could be exploited for potential therapeutic approaches.

## Materials and methods

### Cell culture

*Toxoplasma gondii* tachyzoites were cultured in human foreskin fibroblasts (HFF), sourced from ATCC (SCRC-1041). HFFs and parasites were cultured in Dulbecco’s Modified Eagle’s Medium (DMEM), containing 4.5 gL^-1^ glucose, supplemented with 10% (v/v) fetal bovine serum, 4 mM L-glutamine and penicillin/streptomycin and gentamycin antibiotics and grown at 37°C with 5% CO_2_. When needed anhydrotetracycline (ATc) was added to the medium at a final concertation of 0.5 µM.

### Parasite Genetic manipulation

Parasite genetic manipulation was performed as described previously (31, 35). CRISPR guided promoter replacement and C-terminal triple HA epitope tagging was performed in the TatiΔ*ku80* background (28). gRNAs targeting the start or stop codon of the GOI were designed using the ChopChop tool (https://chopchop.cbu.uib.no/) and cloned into a vector containing the U6 promoter and expressing CAS9-GFP (Tub-Cas9-YFP-pU6-ccdB-tracrRNA) (71) using the BsaI restriction site. For promoter replacement, the pDTS4myc plasmid (28) was used as a template for amplification of DHFR selectable cassette and ATc repressible promoter by PCR. For HA epitope tagging, the CAT selection and triple HA epitope tag were amplified by PCR from p3HA.LIC.CATΔpac plasmid (28, 72). The gRNA/CAS9 vector-PCR product mixture was transfected into the relevant parental line by electroporation and cassette integration selected by antibiotics. Positive clones were isolated by serial dilution and confirmed by PCR analysis, using the primers from Table S3.

Down-regulation of transcript in the promoter replacement line was confirmed by qRT-PCR as described previously (31, 35).

For the complementation of the cKD-ABCB7-HA cell line, ABCB7 sequence with added MfeI and NsiI restriction sites was synthesised (GenScript) and cloned into a pTUB8mycGFPMyoATy expression vector (38). The vectors were electroporated into cKD-ABCB7-HA and integration selected with mycophenolic acid (25 mgmL^-1^) and xanthine (50 mgmL^-1^).

### Growth analysis

Plaque assay: HFF monolayers infected with freshly egressed tachyzoites and grown in the presence or absence of ATc for 8 days. Cells were fixed with methanol and stained with a 0.4% crystal violet solution. 50 plaques per condition were measured from three biological replicates using Image J.

Replication assay: parental and cKD-ABCB7-HA parasites were grown in the presence or absence of ATc for two days and then allowed to infect a fresh HFF monolayer for another day in the same conditions. Immunofluorescence assays were performed as described below using the GAP45 antibody (73). The number of vacuoles containing one, two, four or eight and more parasites were counted for over 100 vacuoles per line per experiment. Three independent experiments were performed.

### SDS and Native PAGE and immunoblot analysis

SDS-PAGE analysis was performed as described previously (31). Immunoblot analysis after SDS-PAGE was performed with relevant primary antibodies: anti-HA (1:500, anti-rat, Sigma), anti-TOM40 (1:2000, anti-rabbit, (34)), anti-Ty (1:1000, anti-mouse, (74)), anti-ATPβ (1:2000, anti-rabbit, Agrisera AS05 085) and anti-CDPK1 (1:10,000, anti-guinea pig, (75)). Immunolabelling was quantified using Odyssey Image Studio (Version 5.2) software, and each HA sample normalised to the TOM40 loading control.

For BN-PAGE parasites were resuspended in solubilisation buffer (750 mM aminocaproic acid, 50 mM Bis-Tris-HCl pH7.0, 0.5 mM EDTA, 1% (w/v) βDDM or SDS) and incubated on ice for 30 minutes. The samples were then centrifuged at 16,000 x g at 4°C for 30 minutes. For βDDM-solubilised samples, the supernatant was combined with sample buffer containing Coomassie G250, resulting in a final concentration of 0.25% DDM and 0.0625% Coomassie G250. For SDS solubilised samples, the β-mercaptoethanol was added and the sample heated at 95°C for 5 minutes before adding sample buffer containing G250. Samples were separated on a NativePAGE 4-16% Bis-Tris gel and transferred to PVDF membrane (0.45 μm, Hybond) using wet transfer in Towbin buffer (25 mM TRIS, 192 mM Glycine, 10% Methanol) for 60 minutes at 100 V, before immunoblot analysis. Bovine mitochondrial membranes were used a molecular weight marker.

### Immunofluorescence assay

To assess localisation immunofluorescence assays were performed as described previously (31) using anti-HA (1:500, anti-rat,Sigma), anti-MYS (1:1000, anti-rabbit, (32)), anti-Ty (1:800, anti-mouse, (74)) and anti-GAP45 (1:1000, anti-rabbit, (73)) primary antibodies. Images were acquired as Z-stacks on a DeltaVision Core microscope (Applied Precision) using the 100x objective and images were processed and deconvolved using the SoftWoRx and FIJI software.

To assess mitochondrial morphology parasites were grown in an HFF monolayer on a glass coverslip in the presence or absence of ATc for three days and an immunofluorescence assay performed using primary antibodies against the mitochondrial protein TOM40 (1:1000, anti-rabbit, (34)). Mitochondrial morphology for each vacuole was scored as “normal” or “abnormal” based on the presence of distinctive lasso-shaped mitochondrial morphology (32). Four independent experiments were performed and over 150 vacuoles were scored for each replicate.

### Respiratory measurements

Extracellular flux analysis: basal oxygen consumption rate (OCR) and extracellular acidification rate (ECAR) was measured using a Seahorse XF HS Mini Analyser (Agilent technologies) as described previously (35).

### Puromycin incorporation assay

The puromycin incorporation assay to assess protein translation was adapted from (45). Parasites were grown in the presence or absence of ATc for three days. Freshly egressed parasites were then filtered through a 3.0 µm polycarbonate filter to remove host cell debris and washed once with media. Parasites were then incubated with media containing 10 µgml^-1^ puromycin (Puromycin dihydrochloride from *Streptomyces alboniger*, Sigma) for 15 minutes at 37 °C, before washing with ice cold PBS. As a control, parasites were incubated with 100 µgml^-1^ cycloheximide for 10 minutes prior to puromycin treatment. Whole cell samples were then analysed by immunoblot using anti-puromycin antibody (1:2000, anti-mouse, clone 12D10, Merck). As a loading control a duplicate SDS-PAGE was performed, and gels incubated in InstantBlue Coomassie protein stain (abcam) for 1 hour to visualise total protein. Puromycin labelling was quantified using Odyssey Image Studio (Version 5.2) software. Total protein was quantified by densitometry analysis using FIJI software. For immunofluorescence-based assay parasites were grown in an HFF monolayer on a glass coverslip in the presence of ATc for three days and then incubated with media containing 10 µgml^-1^ puromycin for 10 minutes at 37 °C. Cells were then fixed and immunofluorescence assay carried out as described above with anti-puromycin antibody at a concentration of 1:1000.

### Immunoprecipitation

For immunoprecipitation of cKD-ABCB7-HA/ ABCB7-Ty, 1.5 x 10^8^ cells were divided into three equal aliquots and incubated in lysis buffer (1% digitonin, 50 mM Tris-HCL pH 7.4, 150 mM NaCl, 2 mM EDTA) for 10 minutes on ice then 30 minutes on a rotator at 4 °C. One aliquot was retained as an input sample while the other two were cleared by centrifugation at 18,000 x g at 4 °C for 30 minutes and the supernatant incubated with either anti-HA or anti-Ty agarose beads overnight at 4°C. The agarose beads were pelleted, and the supernatant was TCA precipitated to give the “unbound” fraction. The beads were then washed in lysis buffer with 0.1% digitonin and resuspended in laemmli buffer to give the “bound” fraction. All fractions were then subjected to immunoblot analysis with antibodies against HA, Ty, TOM40 and CDPK1, as above.

### Sodium carbonate extractions

Sodium carbonate extractions were performed as described previously (35). Briefly, parasites were treated with 100 mM Na_2_CO_3_ pH 11.5 and incubated at 4 °C for 2 hours. The pellet, containing integral membrane proteins, and supernatant, containing peripheral membrane and non-membrane associated proteins, were separated by ultracentrifugation at 189,000 x g. To test the solubility of proteins, parasites were resuspended in 1% triton X-100, incubated at 4 °C for 2 hours, and pellet and supernatant fractions separated by centrifugation at 16,000 x g. Fractions were then tested by immunoblot analysis.

### Cyst quantification

For cyst labelling, intracellular parasites grown in the presence of ATc were processed as above for IFA. Samples were incubated with a biotinylated version of the lectin from *Dolichos biflorus* (Sigma Aldrich, #L6533-5MG) at 1:300, followed by incubation with a Streptavidin-Fluorescein Isothiocyanate (FITC) Conjugate (ThermoFisher #SA10002) at 1:500. Parasites were co-stained with the rabbit anti-IMC3 antibody (76) at 1:1000. At least 100 cysts/vacuoles were counted for each condition in randomly selected fields under a Zeiss AXIO Imager Z2 epifluorescence microscope of the Montpellier Ressources Imagerie platform, driven by the ZEN software v2.3 (Zeiss).

### Quantitative label-free mass spectrometry

The analysis was performed as described previously (21). Parasites of the TATiΔ*ku80* and cKD-ABCB7-HA cell lines were grown for three days in the presence of ATc before being mechanically released from their host cells, filtered on a glass wool fibre column, washed in Hanks’ Balanced Salt Solution (Gibco). After parasite pellet resuspension in SDS lysis buffer (50 mm Tris-HCl pH 8, 10 mm EDTA pH 8, 1% SDS), protein quantification was performed with a bicinchoninic acid assay kit (Abcam) and for each condition, 20 µg of total proteins were separated on a 12% SDS-PAGE run for 20 min at 100 V, stained with colloidal blue (Thermo Fisher Scientific). Each lane was cut in three identical fractions and trypsin digestion and peptide extraction was performed as described previously (77). The LC-MS/MS experiments were performed with an Ultimate 3000 RSLC nano system (Thermo Fisher Scientific Inc, Waltham, MA, USA) interfaced online with a nano easy ion source and an Exploris 240 Plus Orbitrap mass spectrometer (Thermo Fisher Scientific Inc, Waltham, MA, USA).

The .raw files were analysed with MaxQuant version 2.0.3.0 using default settings (78). The minimal peptide length was set to 6. Up to two missed cleavages were allowed. The mass tolerance for the precursor was 20 and 4.5 ppm for the first and the main searches respectively, and for the fragment ions was 20 ppm. The files were searched against *T. gondii* proteome (March 2020 -https://www.uniprot.org/proteomes/UP000005641-8450 entries). Identified proteins were filtered according to the following criteria: at least two different trypsin peptides with at least one unique peptide, an E value below 0.01 and a protein E value smaller than 0.01 were required. Using the above criteria, the rate of false peptide sequence assignment and false protein identification were lower than 1%. Proteins were quantified by label-free method with MaxQuant software using unique and razor peptides intensities (79). Statistical analyses were carried out using RStudio package software. The protein intensity ratio (protein intensity in mutant/protein intensity in parent) and statistical tests were applied to identify the significant differences in the protein abundance. Hits were retained if they were quantified in at least three of the four replicates in at least one experiment. Proteins with a significant (p < 0.05 with Benjamini correction) change in quantitative ratio were considered as significantly up-regulated and down-regulated respectively. Additional candidates that consistently showed absence or presence of LFQ values versus the control in at least 3 out of the four biological replicates were also considered.

### Data availability

All raw MS data and MaxQuant files generated have been deposited to the ProteomeXchange Consortium via the PRIDE partner repository (https://www.ebi.ac.uk/pride/archive) with the dataset identifier PXD048386.

## Supporting information

Table S1: Summary of label-free quantitative proteomics results.

Table S2: Proteins of increased or decreased abundance upon depletion of ABCB7.

Table S3: Summary of oligonucleotides used in this study.

## Acknowledgements

We would like to than Leandro Lemgruber from the imaging facilities of the Wellcome Centre for Integrative Parasitology. We would like to thank Blythe Argyle for assistance with the replication and mitochondrial morphology assays. Mass spectrometry experiments were carried out using the facilities of the Montpellier Proteomics Platform (PPM, MSPP site, BioCampus Montpellier). We thank EupathDB and ToxoDB for providing useful and free access to genome databases. This work was supported by a grant from the Agence Nationale de la Recherche (to S. Besteiro, ANR-22-CE20-0026); by a Early Career Award from the Wellcome Trust (225677/Z/22/Z) (to M. A. Sloan); by a Wellcome Investigator Award (217173_Z_19_Z) (to L. Sheiner) and by a FutureScope award from the Wellcome Centre for Integrative Parasitology and Sir Henry Wellcome Fellowship (to A. E. Maclean, grant number 221681/Z/20/Z).

## Supplementary Figures

**Figure S1.**
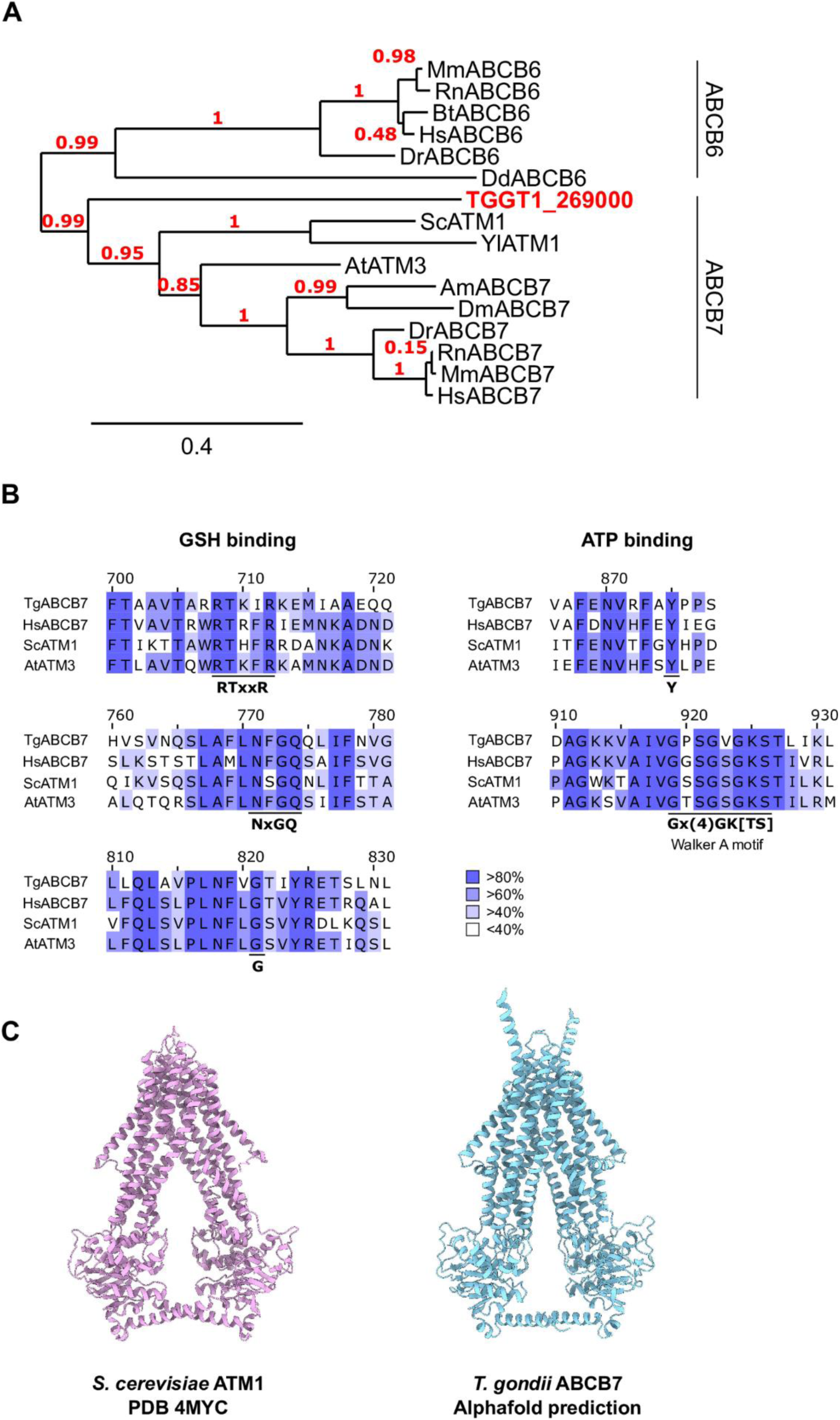
Alignments and phylogenetic tree of ABCB7 homologs. (A) Phylogenetic tree of TGGT1_269000 with ABCB6 and ABCB7 homologs from various species (*Mm*: *Mus musculus*; *Rn*: *Rattus norvegicus*; *Bt*: *Bos taurus*; *Hs*: *Homo sapiens*; *Dr*: *Danio rerio*; *Dd*: *Dictyostelium discoideum*; *Sc*: *Saccharomyces cerevisiae*; *Yl*: *Yarrowia lipolytica*; *At*: *Arabidopsis thaliana*; *Am*: *Apis mellifera*; *Dm*: *Drosophila melanogaster*). Sequences were aligned by MUSCLE and the tree built by PhylML, using the “one-click” mode on phylogeny.fr (80). Values in red are branch support values. (B) Alignments of TGGT1_269000 and HsABCB7 (O75027), ScATM1 (P40416) and AtATM3 (Q9LVM1), showing conservation of key residues involved in glutathione and ATP binding. Alignments were made using Clustal Omega and visualised using JalView. Colour coding depicts percent identity. (C) ATM1 dimer structure from *Saccharomyces cerevisiae* (PDB 4YMC, (14)) and predicted structure of *Toxoplasma gondii* TGGT1_269000 (Alphafold2;(26, 27)). Unstructured loop regions of ABCB7 have been truncated for clarity. Visualised using ChimeraX.

**Figure S2.**
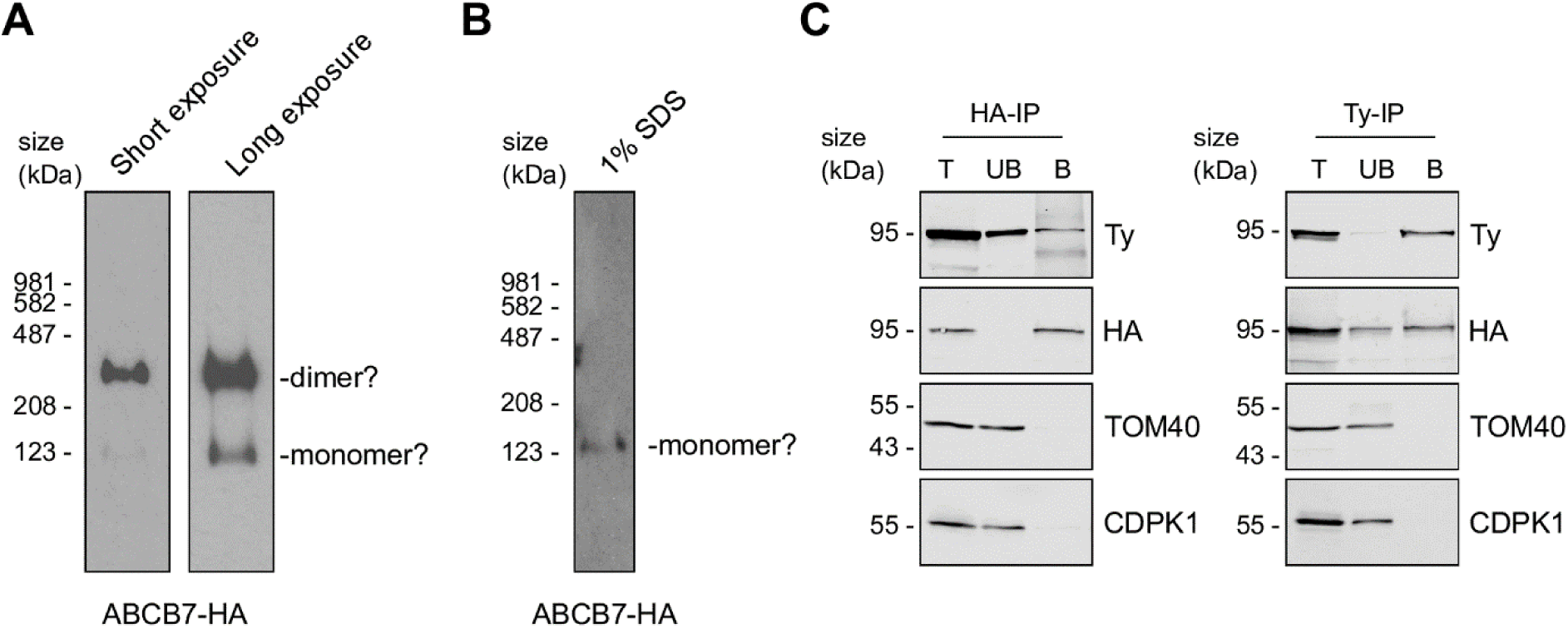
ABCB7 homodimer formation. (A) BN-PAGE analysis of ABCB7-HA parasites extracted in 1% βDDM, immunolabelled with anti-HA. Two exposure lengths shown, and putative monomer and dimer bands indicated. The short exposure is also used in Fig. 1C (B) BN-PAGE analysis of ABCB7-HA parasites extracted in 1% SDS, immunolabelled with anti-HA. (C) Immunoblot analysis of whole cell lysate extracted from cKD-ABCB7-HA + ABCB7-Ty and immunoprecipitated with anti-HA or anti-Ty antibody coupled beads, to produce total lysate (T), unbound (UB) and bound (B) fractions. Samples were separated by SDS-PAGE, blotted, and detected using anti-HA and anti-Ty antibodies to label immunoprecipitated proteins, and anti-TOM40 as an unrelated mitochondrial protein control.

**Figure S3.**
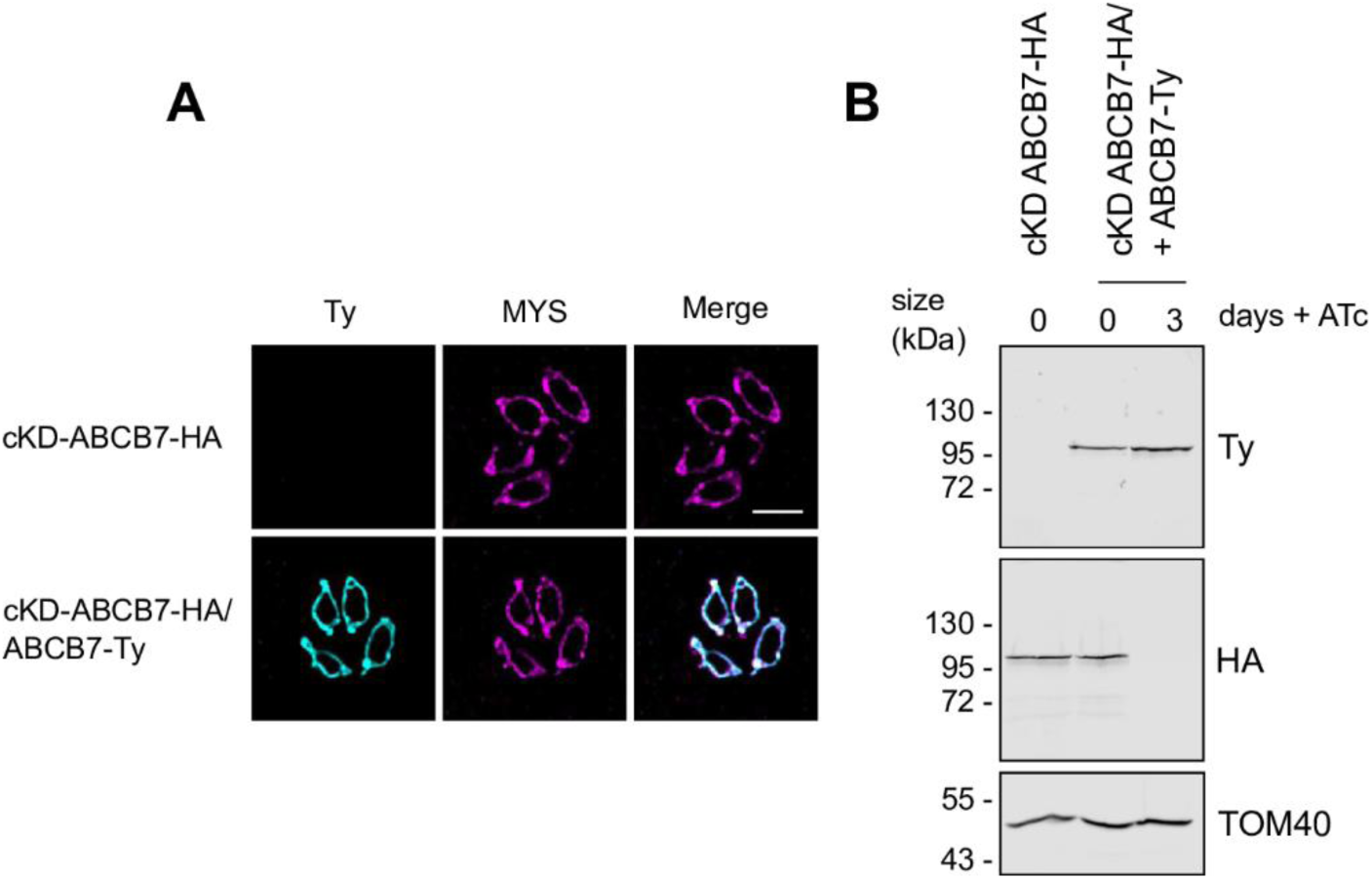
Validation of the complementation of cKD-ABCB7-HA. (A) Immunofluorescence assay analysis of cKD-ABCB7-HA and cKD-ABCB7-HA + ABCB7-Ty parasites, labelled with anti-Ty to detect ABCB7-Ty (cyan), and the mitochondrial marker protein MYS (magenta). Scale bar is 5 µM. (B) Immunoblot analysis of whole cell lysate extracted from cKD-ABCB7-HA and cKD-ABCB7-HA + ABCB7-Ty parasites treated with ATc for zero or three days. Samples were separated by SDS-PAGE, blotted, and detected using anti-Ty, to visualise ABCB7-Ty, anti-HA, to visualise ABCB7-HA, and anti-TOM40 as a loading control.

**Figure S4.**
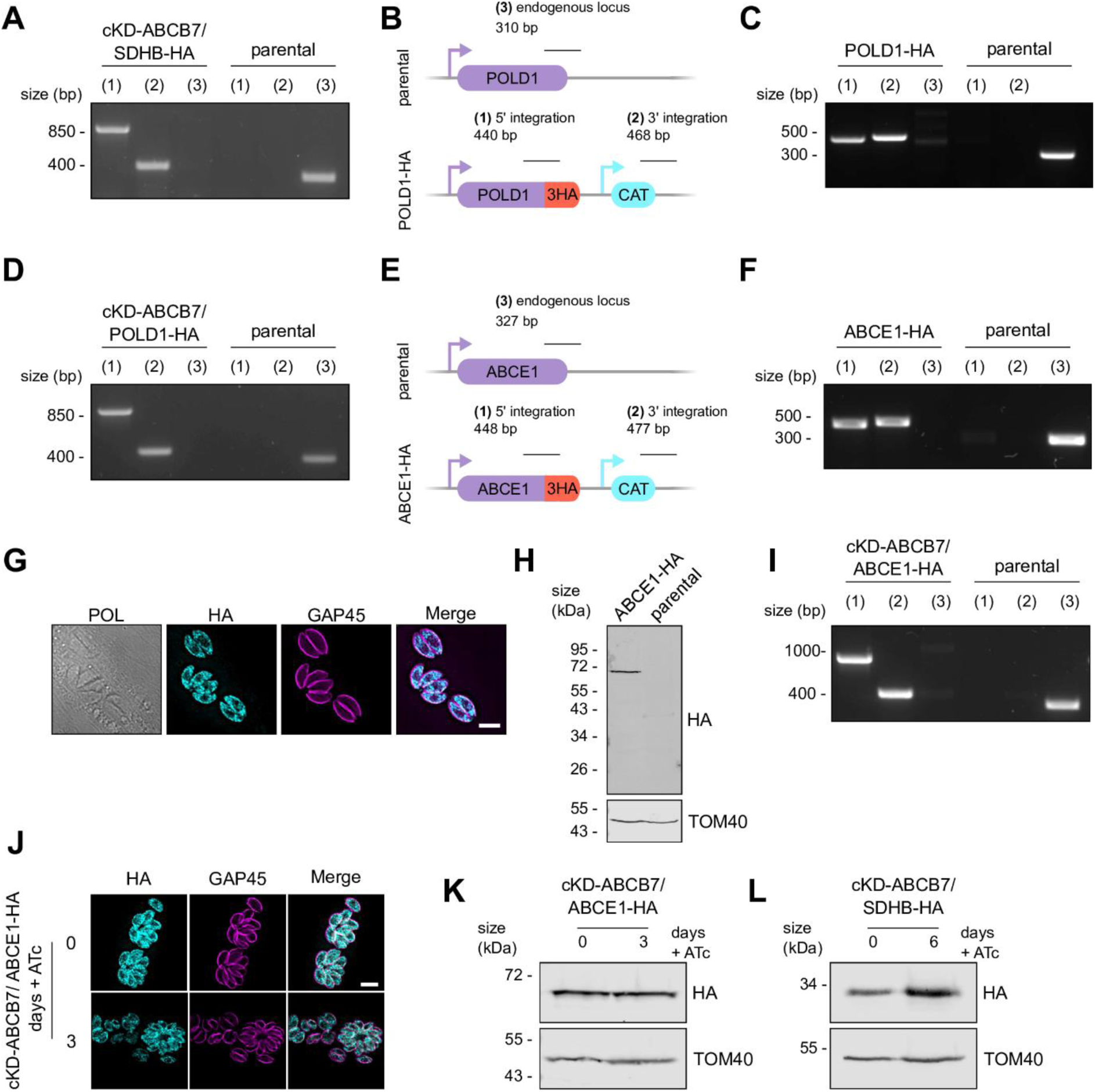
Validation of the genetic manipulation performed for each of the described parasite lines. (A) PCR to test the integration of the DHFR selection cassette and the regulatable promoter into the endogenous locus of SDHB, as outlined in Fig. 2A. (B) Schematic of the strategy used to C-terminally HA-epitope tag the POLD1 protein. The expected size of integration PCRs are shown. (C) PCR to test integration of the HA-epitope tag and CAT selection cassette into the endogenous locus of POLD1, as outlined in *(B)*. (D) PCR to test integration of DHFR selection cassette and the regulatable promoter into the endogenous locus of POLD1, as outlined in Fig. 2A. (E) Schematic of the strategy used to C-terminally HA-epitope tag the ABCE1 protein. The expected size of integration PCRs are shown. (F) PCR to test the integration of the HA-epitope tag and CAT selection cassette into the endogenous locus of ABCE1, as outlined in *(E)*. (G) Immunofluorescence assay analysis of ABCE1-HA parasites, labelled with anti-HA to detect ABCE1-HA (cyan), and the GAP45 (magenta). Scale bar is 5 µM. (H) Immunoblot analysis of whole cell lysate extracted from ABCE1-HA and parental parasites. Samples were separated by SDS-PAGE, blotted, and detected using anti-HA, to visualise ABCE1-HA, and anti-TOM40 as a loading control. (I) PCR to test integration of DHFR selection cassette and the regulatable promoter into the endogenous locus of ABCE1, as outlined in Fig. 2A. (J) Immunofluorescence assay analysis of cKD-ABCB7/ ABCE1-HA parasites, grown in the presence or absence of ATc for three days, labelled with anti-HA to detect ABCE1-HA (cyan), anti-GAP45 to outline the parasites (magenta). Scale bar is 5 µM. (K) Immunoblot analysis of whole cell lysate extracted from cKD-ABCB7/ ABCE1-HA and grown in the presence or absence of ATc for three days. (L) Immunoblot analysis of whole cell lysate extracted from cKD-ABCB7/ SDHB-HA and grown in the presence or absence of ATc for six days.

**S5 Fig.**
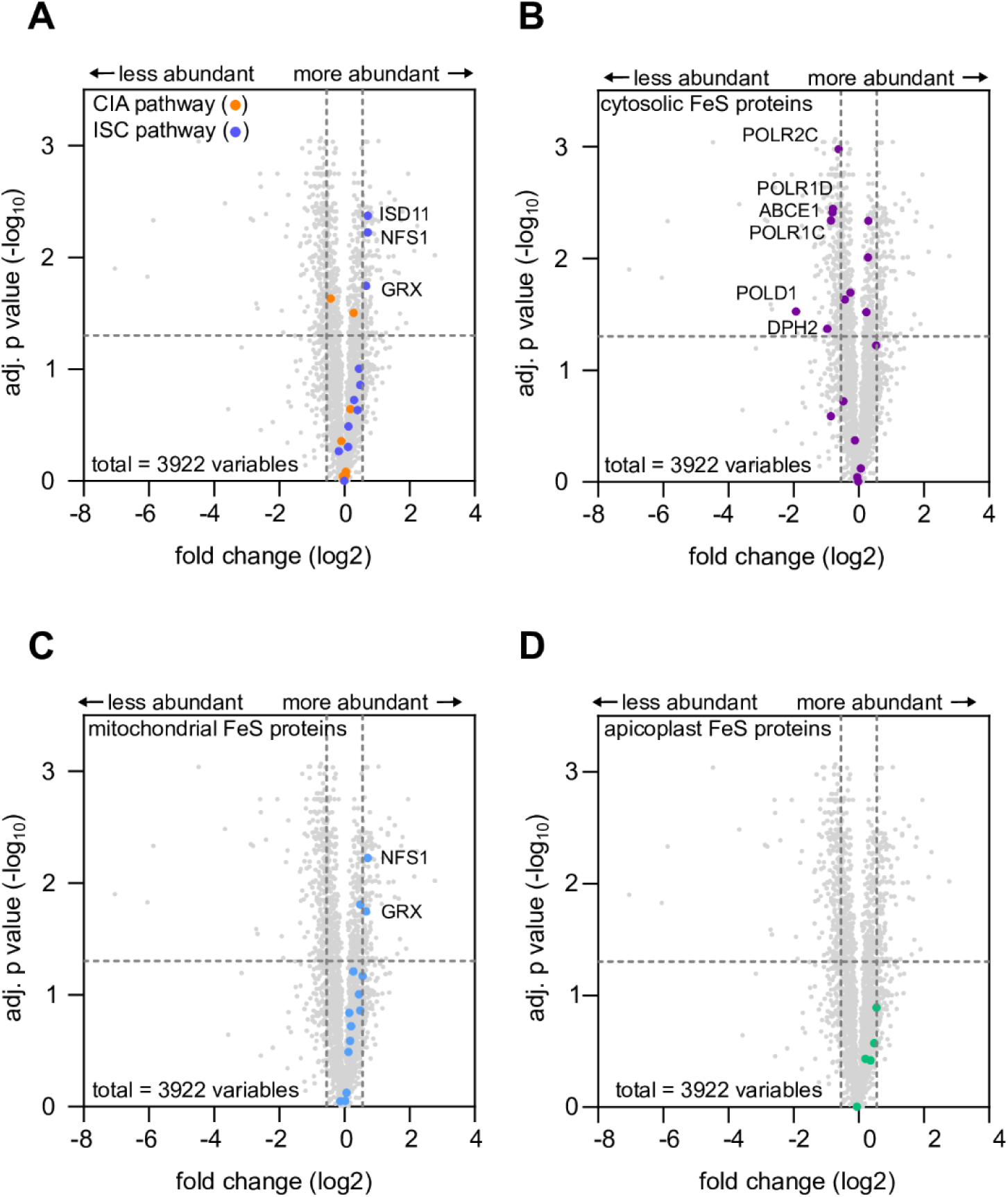
Volcano plots of CIA and ISC pathway proteins and cytosolic, mitochondrial and apicoplast Fe-S proteins in the LFQ proteomic dataset. Volcano plot showing the difference in protein abundance in cKD-ABCB7, as in Fig. 5A, for CIA and ISC pathway member (A), for predicted cytosolic (B), mitochondrial (C) and apicoplast localised Fe-S proteins (see Table S1,2). Proteins that are significantly different from the parental control, and above the ± ≥ 0.55 Log2FC cut-off are individually labelled.

## Supplementary Tables

**Table S1:** Summary of label-free quantitative proteomics results.

**Table S2:** Proteins of increased or decreased abundance upon depletion of ABCB7.

**Table S3:** Summary of oligonucleotides used in this study.

## References

1. Lill R. 2009. Function and biogenesis of iron-sulphur proteins. Nature 460:831–8.

2. Netz DJA, Mascarenhas J, Stehling O, Pierik AJ, Lill R. 2014. Maturation of cytosolic and nuclear iron-sulfur proteins. Trends Cell Biol 24:303–12.

3. Braymer JJ, Freibert SA, Rakwalska-Bange M, Lill R. 2021. Mechanistic concepts of iron-sulfur protein biogenesis in Biology. Biochim Biophys Acta Mol Cell Res 1868:118863.

4. Lill R, Srinivasan V, Mühlenhoff U. 2014. The role of mitochondria in cytosolic-nuclear iron– sulfur protein biogenesis and in cellular iron regulation. Curr Opin Microbiol 22:111–9.

5. Sipos K, Lange H, Fekete Z, Ullmann P, Lill R, Kispal G. 2002. Maturation of cytosolic iron-sulfur proteins requires glutathione. J Biol Chem 277:26944–9.

6. Thomas C, Aller SG, Beis K, Carpenter EP, Chang G, Chen L, Dassa E, Dean M, Duong Van Hoa F, Ekiert D, Ford R, Gaudet R, Gong X, Holland IB, Huang Y, Kahne DK, Kato H, Koronakis V, Koth CM, Lee Y, Lewinson O, Lill R, Martinoia E, Murakami S, Pinkett HW, Poolman B, Rosenbaum D, Sarkadi B, Schmitt L, Schneider E, Shi Y, Shyng S-L, Slotboom DJ, Tajkhorshid E, Tieleman DP, Ueda K, Váradi A, Wen P-C, Yan N, Zhang P, Zheng H, Zimmer J, Tampé R. 2020. Structural and functional diversity calls for a new classification of ABC transporters. FEBS Lett 594:3767–3775.

7. Schaedler TA, Faust B, Shintre CA, Carpenter EP, Srinivasan V, van Veen HW, Balk J. 2015. Structures and functions of mitochondrial ABC transporters. Biochem Soc Trans 43:943–51.

8. Kispal G, Csere P, Prohl C, Lill R. 1999. The mitochondrial proteins Atm1p and Nfs1p are essential for biogenesis of cytosolic Fe/S proteins. EMBO J 18:3981–9.

9. Csere P, Lill R, Kispal G. 1998. Identification of a human mitochondrial ABC transporter, the functional orthologue of yeast Atm1p. FEBS Lett 441:266–70.

10. Bernard DG, Cheng Y, Zhao Y, Balk J. 2009. An allelic mutant series of ATM3 reveals its key role in the biogenesis of cytosolic iron-sulfur proteins in Arabidopsis. Plant Physiol 151:590– 602.

11. Li P, Hendricks AL, Wang Y, Villones RLE, Lindkvist-Petersson K, Meloni G, Cowan JA, Wang K, Gourdon P. 2022. Structures of Atm1 provide insight into [2Fe-2S] cluster export from mitochondria. Nat Commun 13:4339.

12. Fan C, Rees DC. 2022. Glutathione binding to the plant AtAtm3 transporter and implications for the conformational coupling of ABC transporters. Elife 11.

13. Ellinghaus TL, Marcellino T, Srinivasan V, Lill R, Kühlbrandt W. 2021. Conformational changes in the yeast mitochondrial ABC transporter Atm1 during the transport cycle. Sci Adv 7:eabk2392.

14. Srinivasan V, Pierik AJ, Lill R. 2014. Crystal structures of nucleotide-free and glutathione-bound mitochondrial ABC transporter Atm1. Science 343:1137–40.

15. Yan Q, Shen Y, Yang X. 2022. Cryo-EM structure of AMP-PNP-bound human mitochondrial ATP-binding cassette transporter ABCB7. J Struct Biol 214:107832.

16. Schaedler TA, Thornton JD, Kruse I, Schwarzländer M, Meyer AJ, van Veen HW, Balk J. 2014. A conserved mitochondrial ATP-binding cassette transporter exports glutathione polysulfide for cytosolic metal cofactor assembly. J Biol Chem 289:23264–74.

17. Li J, Cowan JA. 2015. Glutathione-coordinated [2Fe-2S] cluster: a viable physiological substrate for mitochondrial ABCB7 transport. Chem Commun (Camb) 51:2253–5.

18. Pandey AK, Pain J, Dancis A, Pain D. 2019. Mitochondria export iron-sulfur and sulfur intermediates to the cytoplasm for iron-sulfur cluster assembly and tRNA thiolation in yeast. J Biol Chem 294:9489–9502.

19. Pandey A, Pain J, Dziuba N, Pandey AK, Dancis A, Lindahl PA, Pain D. 2018. Mitochondria Export Sulfur Species Required for Cytosolic tRNA Thiolation. Cell Chem Biol 25:738–748.e3.

20. Ciofi-Baffoni S, Nasta V, Banci L. 2018. Protein networks in the maturation of human iron-sulfur proteins. Metallomics 10:49–72.

21. Pamukcu S, Cerutti A, Bordat Y, Hem S, Rofidal V, Besteiro S. 2021. Differential contribution of two organelles of endosymbiotic origin to iron-sulfur cluster synthesis and overall fitness in Toxoplasma. PLoS Pathog 17:e1010096.

22. Renaud EA, Pamukcu S, Cerutti A, Berry L, Lemaire-Vieille C, Yamaryo-Botté Y, Botté CY, Besteiro S. 2022. Disrupting the plastidic iron-sulfur cluster biogenesis pathway in Toxoplasma gondii has pleiotropic effects irreversibly impacting parasite viability. J Biol Chem 298:102243.

23. Aw YTV, Seidi A, Hayward JA, Lee J, Makota FV, Rug M, van Dooren GG. 2021. A key cytosolic iron-sulfur cluster synthesis protein localizes to the mitochondrion of Toxoplasma gondii. Mol Microbiol 115:968–985.

24. Harb OS, Roos DS. 2020. ToxoDB: Functional Genomics Resource for Toxoplasma and Related Organisms. Methods Mol Biol 2071:27–47.

25. Kloehn J, Harding CR, Soldati-Favre D. 2021. Supply and demand-heme synthesis, salvage and utilization by Apicomplexa. FEBS J 288:382–404.

26. Varadi M, Anyango S, Deshpande M, Nair S, Natassia C, Yordanova G, Yuan D, Stroe O, Wood G, Laydon A, Žídek A, Green T, Tunyasuvunakool K, Petersen S, Jumper J, Clancy E, Green R, Vora A, Lutfi M, Figurnov M, Cowie A, Hobbs N, Kohli P, Kleywegt G, Birney E, Hassabis D, Velankar S. 2022. AlphaFold Protein Structure Database: massively expanding the structural coverage of protein-sequence space with high-accuracy models. Nucleic Acids Res 50:D439–D444.

27. Jumper J, Evans R, Pritzel A, Green T, Figurnov M, Ronneberger O, Tunyasuvunakool K, Bates R, Žídek A, Potapenko A, Bridgland A, Meyer C, Kohl SAA, Ballard AJ, Cowie A, Romera-Paredes B, Nikolov S, Jain R, Adler J, Back T, Petersen S, Reiman D, Clancy E, Zielinski M, Steinegger M, Pacholska M, Berghammer T, Bodenstein S, Silver D, Vinyals O, Senior AW, Kavukcuoglu K, Kohli P, Hassabis D. 2021. Highly accurate protein structure prediction with AlphaFold. Nature 596:583–589.

28. Sheiner L, Demerly JL, Poulsen N, Beatty WL, Lucas O, Behnke MS, White MW, Striepen B. 2011. A systematic screen to discover and analyze apicoplast proteins identifies a conserved and essential protein import factor. PLoS Pathog 7:e1002392.

29. Barylyuk K, Koreny L, Ke H, Butterworth S, Crook OM, Lassadi I, Gupta V, Tromer E, Mourier T, Stevens TJ, Breckels LM, Pain A, Lilley KS, Waller RF. 2020. A Comprehensive Subcellular Atlas of the Toxoplasma Proteome via hyperLOPIT Provides Spatial Context for Protein Functions. Cell Host Microbe 28:752–766.e9.

30. Seidi A, Muellner-Wong LS, Rajendran E, Tjhin ET, Dagley LF, Aw VY, Faou P, Webb AI, Tonkin CJ, van Dooren GG. 2018. Elucidating the mitochondrial proteome of Toxoplasma gondii reveals the presence of a divergent cytochrome c oxidase. Elife 7.

31. Maclean AE, Bridges HR, Silva MF, Ding S, Ovciarikova J, Hirst J, Sheiner L. 2021. Complexome profile of Toxoplasma gondii mitochondria identifies divergent subunits of respiratory chain complexes including new subunits of cytochrome bc1 complex. PLoS Pathog 17:e1009301.

32. Ovciarikova J, Lemgruber L, Stilger KL, Sullivan WJ, Sheiner L. 2017. Mitochondrial behaviour throughout the lytic cycle of Toxoplasma gondii. Sci Rep 7:42746.

33. Krogh A, Larsson B, von Heijne G, Sonnhammer EL. 2001. Predicting transmembrane protein topology with a hidden Markov model: application to complete genomes. J Mol Biol 305:567– 80.

34. van Dooren GG, Yeoh LM, Striepen B, McFadden GI. 2016. The Import of Proteins into the Mitochondrion of Toxoplasma gondii. J Biol Chem 291:19335–50.

35. Silva MF, Douglas K, Sandalli S, Maclean AE, Sheiner L. 2023. Functional and biochemical characterization of the Toxoplasma gondii succinate dehydrogenase complex. PLoS Pathog 19:e1011867.

36. Sidik SM, Huet D, Ganesan SM, Huynh M-H, Wang T, Nasamu AS, Thiru P, Saeij JPJ, Carruthers VB, Niles JC, Lourido S. 2016. A Genome-wide CRISPR Screen in Toxoplasma Identifies Essential Apicomplexan Genes. Cell 166:1423–1435.e12.

37. Lacombe A, Maclean AE, Ovciarikova J, Tottey J, Mühleip A, Fernandes P, Sheiner L. 2019. Identification of the Toxoplasma gondii mitochondrial ribosome, and characterisation of a protein essential for mitochondrial translation. Mol Microbiol 112:1235–1252.

38. Herm-Götz A, Weiss S, Stratmann R, Fujita-Becker S, Ruff C, Meyhöfer E, Soldati T, Manstein DJ, Geeves MA, Soldati D. 2002. Toxoplasma gondii myosin A and its light chain: a fast, single-headed, plus-end-directed motor. EMBO J 21:2149–58.

39. Tomita T, Bzik DJ, Ma YF, Fox BA, Markillie LM, Taylor RC, Kim K, Weiss LM. 2013. The Toxoplasma gondii cyst wall protein CST1 is critical for cyst wall integrity and promotes bradyzoite persistence. PLoS Pathog 9:e1003823.

40. Mallo N, Ovciarikova J, Martins-Duarte ES, Baehr SC, Biddau M, Wilde M-L, Uboldi AD, Lemgruber L, Tonkin CJ, Wideman JG, Harding CR, Sheiner L. 2021. Depletion of a Toxoplasma porin leads to defects in mitochondrial morphology and contacts with the endoplasmic reticulum. J Cell Sci 134.

41. Hayward JA, Rajendran E, Zwahlen SM, Faou P, van Dooren GG. 2021. Divergent features of the coenzyme Q:cytochrome c oxidoreductase complex in Toxoplasma gondii parasites. PLoS Pathog 17:e1009211.

42. Leonard RA, Tian Y, Tan F, van Dooren GG, Hayward JA. 2023. An essential role for an Fe-S cluster protein in the cytochrome c oxidase complex of Toxoplasma parasites. PLoS Pathog 19:e1011430.

43. Hayward JA, Rajendran E, Makota FV, Bassett BJ, Devoy M, Neeman T, van Dooren GG. 2022. Real-Time Analysis of Mitochondrial Electron Transport Chain Function in Toxoplasma gondii Parasites Using a Seahorse XFe96 Extracellular Flux Analyzer. Bio Protoc 12:e4288.

44. Valasatava Y, Rosato A, Banci L, Andreini C. 2016. MetalPredator: a web server to predict iron-sulfur cluster binding proteomes. Bioinformatics 32:2850–2.

45. Padgett LR, Lentini JM, Holmes MJ, Stilger KL, Fu D, Sullivan WJ. 2018. Elp3 and RlmN: A tale of two mitochondrial tail-anchored radical SAM enzymes in Toxoplasma gondii. PLoS One 13:e0189688.

46. Stilger KL, Sullivan WJ. 2013. Elongator protein 3 (Elp3) lysine acetyltransferase is a tail-anchored mitochondrial protein in Toxoplasma gondii. J Biol Chem 288:25318–25329.

47. Ast T, Itoh Y, Sadre S, McCoy JG, Namkoong G, Wengrod JC, Chicherin I, Joshi PR, Kamenski P, Suess DLM, Amunts A, Mootha VK. 2024. METTL17 is an Fe-S cluster checkpoint for mitochondrial translation. Mol Cell 84:359–374.e8.

48. Paul VD, Lill R. 2015. Biogenesis of cytosolic and nuclear iron-sulfur proteins and their role in genome stability. Biochim Biophys Acta 1853:1528–39.

49. Schmidt EK, Clavarino G, Ceppi M, Pierre P. 2009. SUnSET, a nonradioactive method to monitor protein synthesis. Nat Methods 6:275–7.

50. Hausmann A, Aguilar Netz DJ, Balk J, Pierik AJ, Mühlenhoff U, Lill R. 2005. The eukaryotic P loop NTPase Nbp35: an essential component of the cytosolic and nuclear iron-sulfur protein assembly machinery. Proc Natl Acad Sci U S A 102:3266–71.

51. Tsaousis AD, Gentekaki E, Eme L, Gaston D, Roger AJ. 2014. Evolution of the cytosolic iron-sulfur cluster assembly machinery in Blastocystis species and other microbial eukaryotes. Eukaryot Cell 13:143–53.

52. Grosche C, Diehl A, Rensing SA, Maier UG. 2018. Iron-Sulfur Cluster Biosynthesis in Algae with Complex Plastids. Genome Biol Evol 10:2061–2071.

53. Pyrih J, Žárský V, Fellows JD, Grosche C, Wloga D, Striepen B, Maier UG, Tachezy J. 2021. The iron-sulfur scaffold protein HCF101 unveils the complexity of organellar evolution in SAR, Haptista and Cryptista. BMC Ecol Evol 21:46.

54. Nakai Y, Nakai M, Lill R, Suzuki T, Hayashi H. 2007. Thio modification of yeast cytosolic tRNA is an iron-sulfur protein-dependent pathway. Mol Cell Biol 27:2841–7.

55. Horáková E, Changmai P, Paris Z, Salmon D, Lukeš J. 2015. Simultaneous depletion of Atm and Mdl rebalances cytosolic Fe-S cluster assembly but not heme import into the mitochondrion of Trypanosoma brucei. FEBS J 282:4157–75.

56. Luo D, Bernard DG, Balk J, Hai H, Cui X. 2012. The DUF59 family gene AE7 acts in the cytosolic iron-sulfur cluster assembly pathway to maintain nuclear genome integrity in Arabidopsis. Plant Cell 24:4135–48.

57. Bernard DG, Netz DJA, Lagny TJ, Pierik AJ, Balk J. 2013. Requirements of the cytosolic iron-sulfur cluster assembly pathway in Arabidopsis. Philos Trans R Soc Lond B Biol Sci 368:20120259.

58. Bastow EL, Bych K, Crack JC, Le Brun NE, Balk J. 2017. NBP35 interacts with DRE2 in the maturation of cytosolic iron-sulphur proteins in Arabidopsis thaliana. Plant J 89:590–600.

59. Bak DW, Weerapana E. 2023. Monitoring Fe-S cluster occupancy across the E. coli proteome using chemoproteomics. Nat Chem Biol 19:356–366.

60. Lill R, Freibert S-A. 2020. Mechanisms of Mitochondrial Iron-Sulfur Protein Biogenesis. Annu Rev Biochem 89:471–499.

61. Hassan MA, Vasquez JJ, Guo-Liang C, Meissner M, Nicolai Siegel T. 2017. Comparative ribosome profiling uncovers a dominant role for translational control in Toxoplasma gondii. BMC Genomics 18:961.

62. Holmes MJ, Augusto L da S, Zhang M, Wek RC, Sullivan WJ. 2017. Translational Control in the Latency of Apicomplexan Parasites. Trends Parasitol 33:947–960.

63. Gastens MH, Fischer H-G. 2002. Toxoplasma gondii eukaryotic translation initiation factor 4A associated with tachyzoite virulence is down-regulated in the bradyzoite stage. Int J Parasitol 32:1225–34.

64. Wang F, Holmes MJ, Hong HJ, Thaprawat P, Kannan G, Huynh M-H, Schultz TL, Licon MH, Lourido S, Sullivan WJ, O’Leary SE, Carruthers VB. 2023. Translation initiation factor eIF1.2 promotes Toxoplasma stage conversion by regulating levels of key differentiation factors. bioRxiv 10.1101/2023.11.03.565545.

65. Holmes MJ, Bastos MS, Dey V, Severo V, Wek RC, Sullivan WJ. 2023. mRNA cap-binding protein eIF4E1 is a novel regulator of Toxoplasma gondii latency. bioRxiv 10.1101/2023.10.09.561274.

66. Jedelský PL, Doležal P, Rada P, Pyrih J, Smíd O, Hrdý I, Sedinová M, Marcinčiková M, Voleman L, Perry AJ, Beltrán NC, Lithgow T, Tachezy J. 2011. The minimal proteome in the reduced mitochondrion of the parasitic protist Giardia intestinalis. PLoS One 6:e17285.

67. Leger MM, Kolisko M, Kamikawa R, Stairs CW, Kume K, Čepička I, Silberman JD, Andersson JO, Xu F, Yabuki A, Eme L, Zhang Q, Takishita K, Inagaki Y, Simpson AGB, Hashimoto T, Roger AJ. 2017. Organelles that illuminate the origins of Trichomonas hydrogenosomes and Giardia mitosomes. Nat Ecol Evol 1:0092.

68. Stairs CW, Eme L, Brown MW, Mutsaers C, Susko E, Dellaire G, Soanes DM, van der Giezen M, Roger AJ. 2014. A SUF Fe-S cluster biogenesis system in the mitochondrion-related organelles of the anaerobic protist Pygsuia. Curr Biol 24:1176–86.

69. Karnkowska A, Vacek V, Zubáčová Z, Treitli SC, Petrželková R, Eme L, Novák L, Žárský V, Barlow LD, Herman EK, Soukal P, Hroudová M, Doležal P, Stairs CW, Roger AJ, Eliáš M, Dacks JB, Vlček Č, Hampl V. 2016. A Eukaryote without a Mitochondrial Organelle. Curr Biol 26:1274–84.

70. Vacek V, Novák LVF, Treitli SC, Táborský P, Cepicka I, Kolísko M, Keeling PJ, Hampl V. 2018. Fe-S Cluster Assembly in Oxymonads and Related Protists. Mol Biol Evol 35:2712– 2718.

71. Curt-Varesano A, Braun L, Ranquet C, Hakimi M-A, Bougdour A. 2016. The aspartyl protease TgASP5 mediates the export of the Toxoplasma GRA16 and GRA24 effectors into host cells. Cell Microbiol 18:151–67.

72. Huynh M-H, Carruthers VB. 2009. Tagging of endogenous genes in a Toxoplasma gondii strain lacking Ku80. Eukaryot Cell 8:530–9.

73. Plattner F, Yarovinsky F, Romero S, Didry D, Carlier M-F, Sher A, Soldati-Favre D. 2008. Toxoplasma profilin is essential for host cell invasion and TLR11-dependent induction of an interleukin-12 response. Cell Host Microbe 3:77–87.

74. Bastin P, Bagherzadeh Z, Matthews KR, Gull K. 1996. A novel epitope tag system to study protein targeting and organelle biogenesis in Trypanosoma brucei. Mol Biochem Parasitol 77:235–9.

75. Waldman BS, Schwarz D, Wadsworth MH, Saeij JP, Shalek AK, Lourido S. 2020. Identification of a Master Regulator of Differentiation in Toxoplasma. Cell 180:359–372.e16.

76. Anderson-White BR, Ivey FD, Cheng K, Szatanek T, Lorestani A, Beckers CJ, Ferguson DJP, Sahoo N, Gubbels M-J. 2011. A family of intermediate filament-like proteins is sequentially assembled into the cytoskeleton of Toxoplasma gondii. Cell Microbiol 13:18–31.

77. Berger N, Vignols F, Przybyla-Toscano J, Roland M, Rofidal V, Touraine B, Zienkiewicz K, Couturier J, Feussner I, Santoni V, Rouhier N, Gaymard F, Dubos C. 2020. Identification of client iron-sulfur proteins of the chloroplastic NFU2 transfer protein in Arabidopsis thaliana. J Exp Bot 71:4171–4187.

78. Cox J, Mann M. 2008. MaxQuant enables high peptide identification rates, individualized p.p.b.-range mass accuracies and proteome-wide protein quantification. Nat Biotechnol 26:1367–72.

79. Cox J, Hein MY, Luber CA, Paron I, Nagaraj N, Mann M. 2014. Accurate proteome-wide label-free quantification by delayed normalization and maximal peptide ratio extraction, termed MaxLFQ. Mol Cell Proteomics 13:2513–26.

80. Dereeper A, Guignon V, Blanc G, Audic S, Buffet S, Chevenet F, Dufayard J-F, Guindon S, Lefort V, Lescot M, Claverie J-M, Gascuel O. 2008. Phylogeny.fr: robust phylogenetic analysis for the non-specialist. Nucleic Acids Res 36:W465–9.

